# Stop codon-proximal 3′UTR introns in vertebrates can elicit EJC-dependent Nonsense-Mediated mRNA Decay

**DOI:** 10.1101/677666

**Authors:** Pooja Gangras, Thomas L. Gallagher, Robert D. Patton, Zhongxia Yi, Michael A. Parthun, Kiel T. Tietz, Natalie C. Deans, Ralf Bundschuh, Sharon L. Amacher, Guramrit Singh

## Abstract

The Exon Junction Complex (EJC) regulates many steps in post-transcriptional gene expression and is essential for cellular function and organismal development; however, EJC-regulated genes and genetic pathways during development remain largely unknown. To study EJC function during zebrafish development, we first established that zebrafish EJCs mainly bind ∼24 nucleotides upstream of exon-exon junctions, and are also detected at more distant non-canonical positions. We then generated mutations in two zebrafish EJC core genes, *rbm8a* and *magoh*, and observed that homozygous mutant embryos show paralysis, muscle disorganization, neural cell death, and motor neuron outgrowth defects. Coinciding with developmental defects, mRNAs subjected to Nonsense-Mediated mRNA Decay (NMD) due to translation termination ≥ 50 nts upstream of the last exon-exon junction are upregulated in EJC mutant embryos. Surprisingly, several transcripts containing 3′UTR introns (3′UI) < 50 nts downstream of a stop codon are also upregulated in EJC mutant embryos. These proximal 3′UI-containing transcripts are also upregulated in NMD-compromised zebrafish embryos, cultured human cells, and mouse embryonic stem cells. Loss of function of *foxo3b,* one of the upregulated proximal 3′UI-containing genes, partially rescues EJC mutant motor neuron outgrowth. In addition to *foxo3b*, 166 other genes contain a proximal 3′UI in zebrafish, mouse and humans, and these genes are enriched in nervous system development and RNA binding functions. A proximal 3′UI-containing 3′UTR from one of these genes, *HNRNPD*, is sufficient to reduce steady state transcript levels when fused to a *β-globin* reporter in HeLa cells. Overall, our work shows that genes with stop codon-proximal 3′UIs encode a new class of EJC-regulated NMD targets with critical roles during vertebrate development.

## Introduction

Post-transcriptional control of gene expression is essential for cellular function and plays a crucial role in cell fate decisions and cell differentiation pathways during development. RNA-binding proteins that control post-transcriptional processes are therefore important regulators of development (1, 2). One set of proteins crucial for post-transcriptional control is the Exon Junction Complex (EJC) that assembles ∼24 nucleotides (nts) upstream of exon-exon junctions during pre-mRNA splicing (3–5). The EJC is comprised of three core proteins, Eif4a3, Rbm8a (Y14), and Magoh, which partner with several peripheral factors to regulate pre-mRNA splicing and mRNA export in the nucleus and mRNA localization, translation and nonsense-mediated mRNA decay (NMD) in the cytoplasm (3–5).

The EJC core components were first discovered in *Drosophila* for their role in germ cell specification and embryo patterning (6, 7). More recent findings that mutations in human EJC core protein-encoding genes cause defects in neural and musculoskeletal development have underscored the importance of the EJC during development (8, 9). Interestingly, developmental defects in neural cell types are observed in *Xenopus* embryos and mouse models with reduced EJC core protein levels, suggesting conserved and essential EJC functions in these cells (10–13). In recent years, work using mouse models has illuminated an important role for EJC core components in neural precursor cell proliferation during brain development (14–16). In mice that are conditionally haploinsufficient for any of one the three EJC core components, neural precursor cells exit the cell cycle early and prematurely differentiate, leading to untimely production of excess neurons, which then undergo p53-dependent apoptosis (10,11,16,17). These defects lead to impaired cortical development and microcephaly, a phenotype also linked with *RBM8A* and *EIF4A3* mutations in humans (8, 9). While these advances highlight the critical role of EJC during neural development, much remains to be learned about EJC-regulated developmental gene expression programs and how each of the EJC’s many functions contribute to developmental gene regulation.

In vertebrates, one key function of the EJC is its ability to trigger NMD when present downstream of a terminating ribosome. Current models state that when a ribosome terminates translation ≥ 50 nts upstream of an exon-exon junction, the EJC that remains bound in the 3′-untranslated region (UTR), through its interactions with peripheral proteins UPF3B and UPF2, activates central NMD factor UPF1 to induce NMD (18, 19). Thus, NMD suppresses the expression of aberrant transcripts bearing premature termination codons. Additionally, NMD also regulates expression of normal transcripts that contain NMD-inducing features such as upstream open reading frames (uORF) and 3′UTR introns (3′UIs) (20–22). In the case of such normal transcripts, the ribosome terminates at the normal stop after production of at least one full length polypeptide, but due to the presence of a downstream EJC, the transcript is targeted for decay. Thus, EJC-dependent NMD also acts as a post-transcriptional mechanism to fine-tune protein expression as has been shown for *ARC* mRNA in neuronal synapses (23, 24). Interestingly, 3′UI-bearing transcripts are enriched for neuronal and hematopoietic functions (23), and are expressed in tissue-specific patterns (24), suggesting that 3′UIs may play an important role in regulating tissue-specific developmental programs via EJC-dependent NMD. However, specific mRNAs, genetic pathways, and developmental events that are regulated by 3′UI-dependent NMD remain largely unknown.

In this work, we establish zebrafish as a model to study EJC developmental functions. As expected from work in human cells, we show that in zebrafish embryos the EJC mainly binds ∼24 nts upstream of exon-exon junctions and is also detected on more distant exonic positions. Zebrafish Rbm8a and Magoh are crucial for early zebrafish embryonic development, with muscle and neural lineages being particularly sensitive to the loss of EJC. We find that EJC-dependent NMD is disrupted in *rbm8a* and *magoh* mutant embryos. Strikingly, we uncover a class of genes that contain a stop codon-proximal 3′UI (intron within 50 nts of the stop codon) and are upregulated in zebrafish EJC mutant embryos and *upf1* morphants. We show that loss of *foxo3b*, a proximal 3′UI-containing gene whose transcript and protein levels are elevated in EJC mutants, partially rescues the EJC motor neuron outgrowth defect. Proximal 3′UI-containing genes are also widespread in human and mouse genomes, and are similarly regulated by human and mouse NMD pathways. We identify 167 genes that contain a 3′UI at a stop-codon proximal position in zebrafish, mouse and humans. These genes are enriched for genes encoding RNA-binding proteins and proteins involved in nervous system development. A 3′UTR containing a proximal intron from one of these genes can trigger NMD when linked to a human *β-globin* reporter RNA.

## Results

### EJC composition and deposition is conserved in zebrafish

The three proteins, Eif4a3, Rbm8a, and Magoh, that form the EJC core are highly conserved among multicellular organisms including zebrafish and humans (Fig. S1A). To test if the zebrafish EJC core proteins assemble into a complex similar to that observed in human cells (25–27) and *Drosophila* cells (28), we immunoprecipitated Rbm8a from RNase-treated zebrafish embryo extracts. We find that both Eif4a3 and Magoh, but not a negative control RNA-binding protein HuC, specifically co-immunopreciptate with Rbm8a (Fig. 1A). We and others have previously shown that the EJC primarily binds 24 nts upstream of exon-exon junctions in cultured human cells and adult *Drosophila* (29–33). To test if the EJC binds at a similar position on zebrafish spliced RNAs, we first optimized RNA-immunoprecipitation (RIP) from RNase-treated zebrafish embryo lysates using an Rbm8a antibody (Fig. S1B-C). By performing Rbm8a RIP-Seq under optimized conditions from zebrafish embryo lysates, we obtained three well-correlated biological replicates of Rbm8a-associated RNA fragments (Fig. S1D, table S1). As expected, Rbm8a footprint read densities are significantly higher in exonic regions as compared to intronic regions (Fig. 1B), and on exons from multi-exon genes as compared to those from intron-less genes (Fig. 1C). A meta-exon analysis shows that, like the human EJC, zebrafish Rbm8a footprints cluster around the canonical EJC binding site 24 nts upstream of exon 3′ ends (Fig. 1D). As expected, 5′ and 3′ ends of RIP-Seq reads accumulate upstream and downstream of the −24 nts position, respectively (Fig. 1E). A dramatic reduction in 5′ and 3′ end read counts in a ∼10 nts region around the −24 position reveals the RNA segment that is protected from RNase digestion by Rbm8a-containing EJCs (Fig. 1E). The predominant Rbm8a-occupancy position close to exonic 3′ ends is also evident from the RIP-Seq read distribution on individual exons (Fig. 1F and Fig. S1E). Qualitatively, many canonical EJC sites from highly expressed multi-exon genes show variable Rbm8a binding (Fig. 1F and Fig. S1E). These profiles also show that zebrafish Rbm8a also associates with non-canonical positions away from the −24 position, as observed previously in human cells (Fig. 1C,1F and S1E). Thus, like in humans, the zebrafish EJC is also detected at non-canonical positions. Taken together, we conclude that the EJC core assembly and its deposition pattern is conserved between zebrafish and humans.

**Figure 1.**
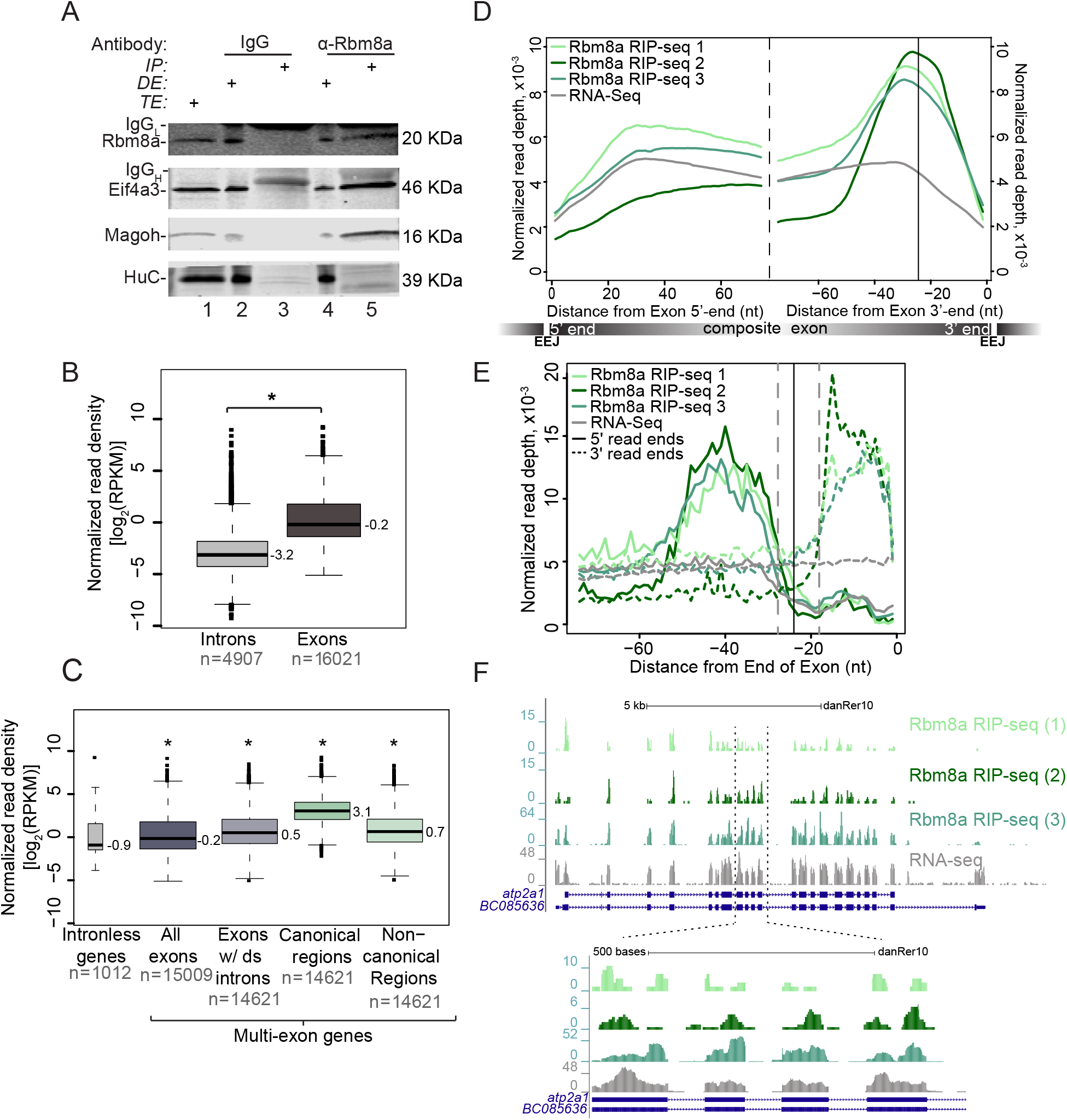
The zebrafish EJC is detected ∼24 nucleotides upstream of exon-exon junctions. A. Western blot indicating Rbm8a, Eif4a3 and Magoh proteins detected in RNase I-treated zebrafish embryo total extract (TE, lane 1), depleted extract (DE, lanes 2 and 4) immunoprecipitated protein complexes (IP, lanes 3 and 5). Antigens detected in the blot are listed on the left and antibodies used to immunoprecipitate complexes are listed on top. The signal corresponding to the antibody light chain and heavy chain in the IP lanes is indicated by IgGL and IgGH respectively. B. Boxplots showing the Rbm8a RIP-Seq normalized read densities (reads per kilobase per million, RPKM) in intronic versus exonic genomic regions. Asterisk at the top indicates Wilcoxon test p-values, which are < 10^-6^. C. Boxplots as in B showing the Rbm8a RIP-Seq normalized read densities (RPKM) in the indicated genomic regions (bottom). Exons with downstream introns include all but last exons. Asterisk at the top indicates Wilcoxon test p-values, which are < 10^-6^. D. Meta-exon plots showing Rbm8a RIP-Seq and RNA-Seq (indicated on the left) normalized read depths in a 75 nt region starting from the exon 5′ (left of dashed black line) or 3′ ends (right of dashed black line). Vertical black line: expected canonical EJC binding site (−24 nt) based on human studies. A composite exon with the relative position of exon-exon junctions (EEJ) is diagrammed at the bottom. E. A meta-exon plot of start and end of Rbm8a RIP-Seq footprint reads (5′ ends, solid lines; 3′ ends, dotted lines). Vertical black line: canonical EJC site (−24 nt). Gray vertical dashed lines represent boundaries of the minimal EJC occupied site. F. Top: UCSC genome browser screenshots showing read coverage along the *atp2a1* gene in the Rbm8a RIP-Seq or RNA-Seq replicates as labeled on the right. Bottom: A zoomed in view of the region between the two dotted lines on the top panel. The y-axis on the left of each track shows maximal read coverage in the shown interval.

### *rbm8a* and *magoh* mutant embryos show defects in motility, muscle organization, and motor neuron outgrowth

To identify the molecular functions of the EJC during embryonic development, we generated zebrafish *rbm8a* and *magoh* mutant embryos. Using a CRISPR/Cas9-based approach (34), we created frame-shifting deletions early in the protein coding sequence to generate null alleles (Fig. 2A). Fish heterozygous for *rbm8a^oz36^* or *magoh^oz37^* alleles displayed no obvious phenotypes and were fully viable. Homozygous mutant *rbm8a* or *magoh* embryos (hereafter collectively referred to as EJC mutant embryos), obtained by intercrossing *rbm8a* or *magoh* heterozygotes, initially appear morphologically normal except for mild head necrosis and tail curvature prior to 24 hpf (hours post fertilization) (Fig. 2B). A closer examination revealed that head necrosis can be readily detected by acridine orange staining at 19 hpf (Fig. S2A) and morphologically by 21 hpf (Fig. S2B). After 24 hpf, EJC mutant embryos decline rapidly, with the decline in *magoh* mutant embryos appearing more advanced at each developmental time point examined (Fig. S2B-C). Both EJC mutant embryos have reduced head size, pericardial edema, and widespread necrosis by 32 hpf, and die by 48 hpf.

**Figure 2.**
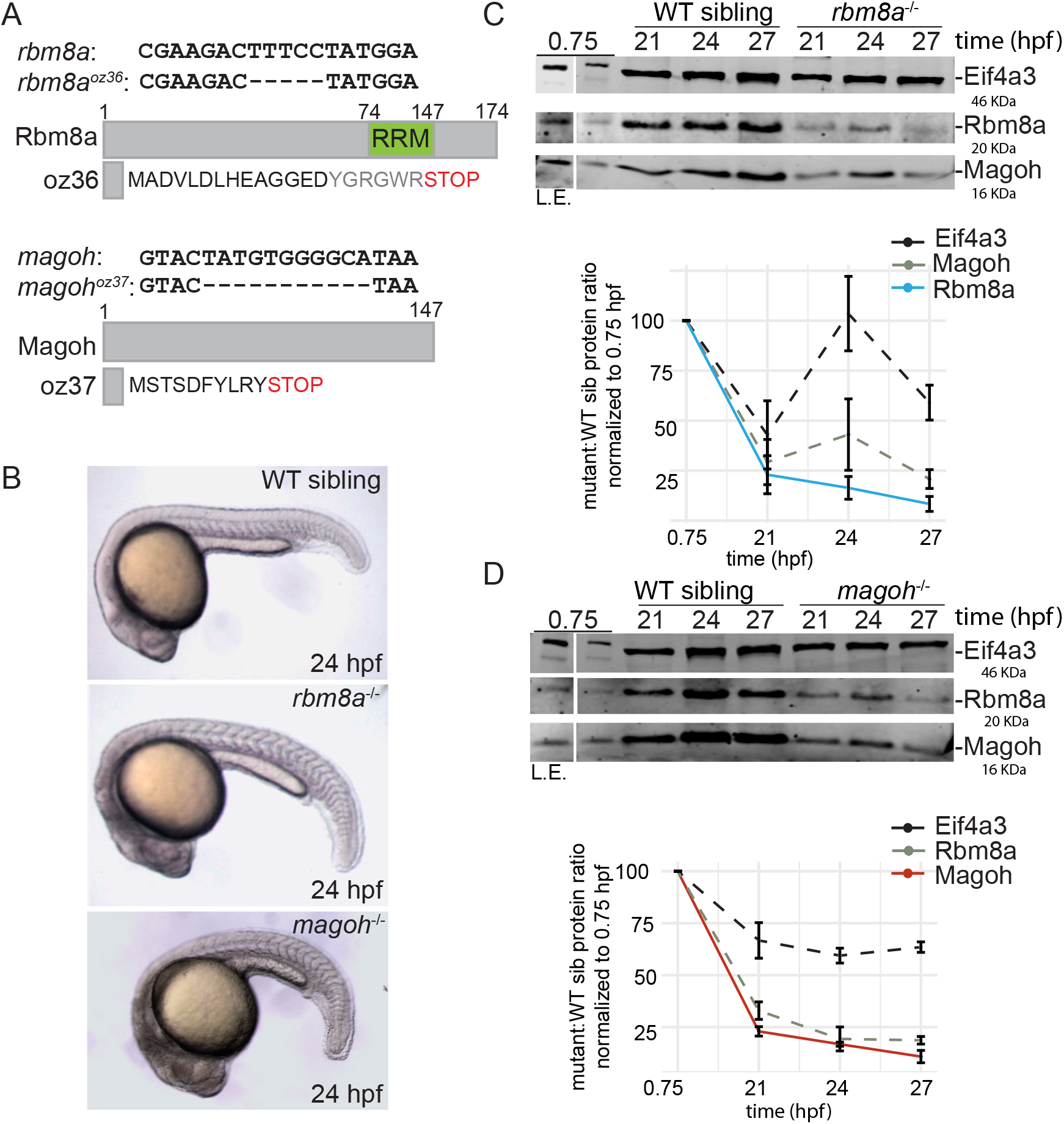
Zebrafish *rbm8a* and *magoh* mutant embryos show gradual loss of maternally contributed Rbm8a and Magoh proteins during early development. A. Schematic illustrating the *rbm8a^oz36^* and *magoh^oz37^* alleles and the predicted proteins they encode. Full length Rbm8a and Magoh proteins are also shown. RRM: RNA Recognition Motif. B. Whole mount images of live wild-type sibling, *rbm8a* mutant, and *magoh* mutant embryos at 24 hpf. Increased grayness in the head region of homozygous *rbm8a* and *magoh* mutant embryos indicates cell death. C. Top: Western blots showing EJC protein expression in wild type (WT) sibling and *rbm8a*^-/-^ mutant embryos. Antigens detected are listed on the right and embryo genotype is listed above the blot. Developmental time points (hpf) are indicated above each lane. Protein from five (0.75 hpf) or ten embryos (all other time points) was loaded in each lane. A longer exposure (L.E.) of the 0.75 hpf lane is on the left. Bottom: Line graphs showing the amount of protein (per embryo) in the mutant embryos compared to wild-type sibling as a percent of protein present at 0.75 hpf. Error bars represent standard error of means. D. Top: Western blots showing EJC protein expression in wild type (WT) sibling and *magoh*^-/-^ mutant embryos. Antigens detected are listed on the right and embryo genotype is listed above the blot. Developmental time points (hpf) are indicated above each lane. Protein from five (0.75 hpf) or ten embryos (all other time points) was loaded in each lane. A longer exposure (L.E.) of the 0.75 hpf lane is on the left. Bottom: Line graphs showing the amount of protein (per embryo) in the mutant embryos compared to wild-type sibling as a percent of protein present at 0.75 hpf. Error bars represent standard error of means.

We hypothesized that EJC mutant embryos are initially sustained by maternally-deposited *rbm8a* and *magoh* transcripts (35) and protein, and that developmental defects appearing at 19-21 hpf coincide with maternal depletion. Consistent with maternal deposition of EJC transcript and/or protein, we detect Rbm8a and Magoh protein in 2-4 cell stage embryos (0.75 hpf) (Fig. 2C-D). Over time, both Rbm8a and Magoh levels in EJC mutant embryos decrease to ∼25% of their respective levels in wild-type siblings, and levels continue to drop over the next six hours (Fig. 2C-D). As previously observed in mammalian cells (36), reduction of either protein of the Rbm8a:Magoh heterodimer leads to a concomitant depletion of the other protein (Fig. 2C-D).

Although EJC mutant embryos are morphologically indistinguishable from wild-type siblings at 18 hpf, we find that they are paralyzed (Fig. 3A). It is unlikely that the lack of spontaneous contractions is due to developmental delay as EJC mutant embryos never become motile (data not shown). To further characterize the paralysis phenotype, we assessed muscle and motor neuron morphology, as these cell types are required for motility. Myosin heavy chain immunostaining reveals that EJC mutant embryos have disorganized myofibers and have U-shaped instead of chevron-shaped myotomes (Fig. 3B-D), with muscle defects in *magoh* mutant embryos consistently more severe than in *rbm8a* mutant embryos. Co-labeling of motor axons (using anti-SV2) and neuromuscular junctions (using Alexa Fluor conjugated α-Bungarotoxin), shows that motor axon length and neuromuscular junction number are reduced in EJC mutant embryos (Fig. 3E-H). Thus, as expected of genes that encode proteins that function as a complex, homozygous *rbm8a* and *magoh* mutant embryos show phenotypically similar muscle organization and motor neuron outgrowth defects.

**Figure 3.**
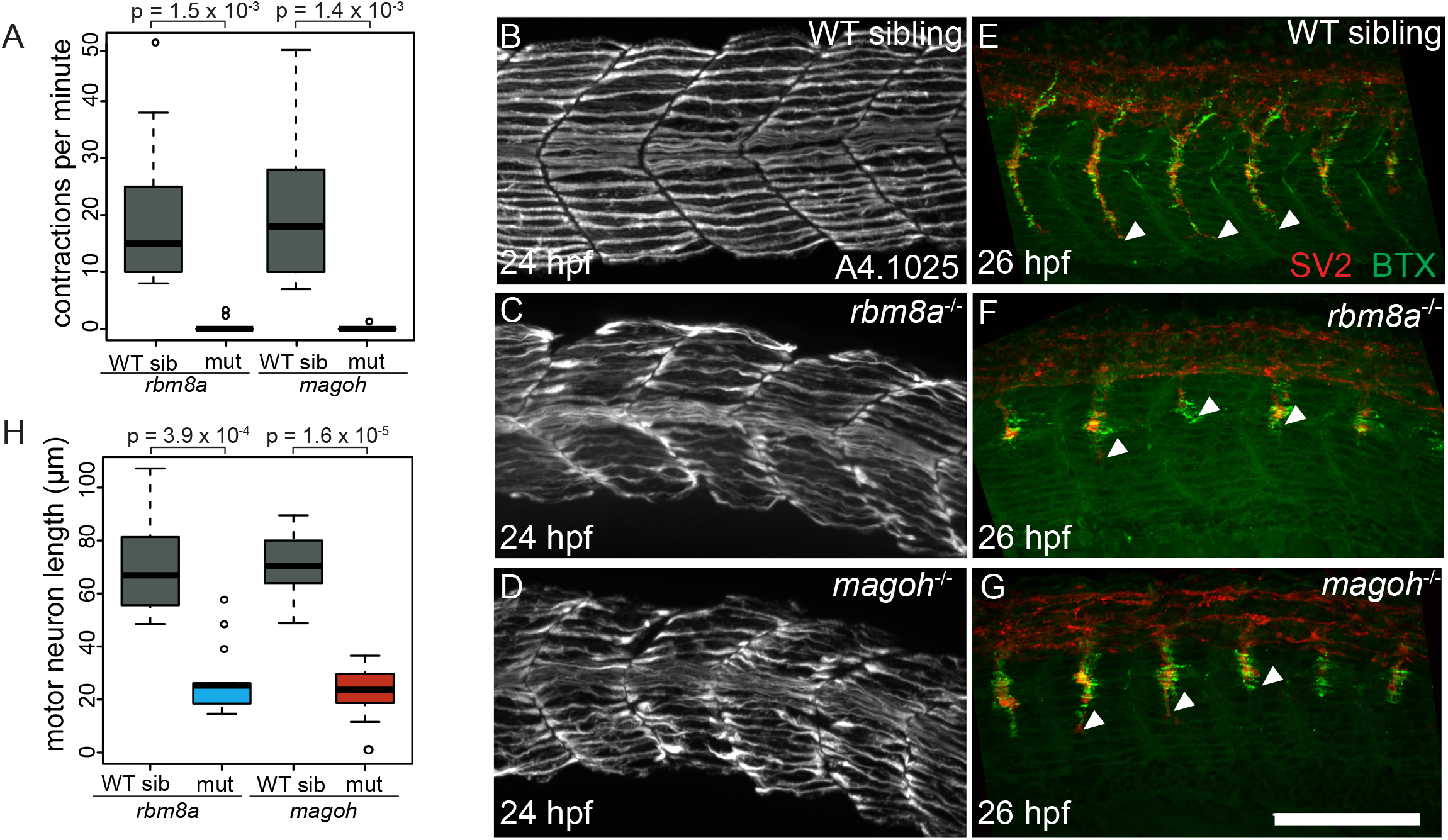
EJC mutant embryos are paralyzed, have disorganized muscles and stunted motor neurons. A. Boxplots showing the number of spontaneous contractions per minute measured for the EJC mutant embryos and WT siblings at 24 hpf as indicated on the x-axis. Welch t-test p-values are indicated at the top. B-D. Immunofluorescence images showing Myh1 expression in somites 10-14 of WT sibling (B) *rbm8a* mutant (C) and *magoh* (D) mutant embryos. Antibody used was anti-A4.1025 (see methods) (N = 10 embryos/genotype). E-G. Merged confocal images of somites 12-16 in WT siblings (E) *rbm8a* (F) and *magoh* (G) mutant embryos showing immunofluorescence detection of motor neurons (anti-SV2; red) and acetylcholine receptors (α-Bungarotoxin; green). Neuro-muscular junctions in the merged image appear yellow. White arrowheads point to the end of the motor neuron. Scale bar in G is 100 nm. H. Boxplots showing the quantification of motor neuron length in somites 12-15 of wild-type sibling, *rbm8a* mutant, and *magoh* mutant embryos (N = 4 embryos/genotype and 4 neurons/embryo). Welch t-test p-values are at the top.

### Analysis of gene expression in *rbm8a* and *magoh* mutant embryos

To identify EJC-regulated genes during zebrafish embryonic development, we performed RNA-Seq of EJC mutant embryos and their wild-type siblings at two developmental time points, 21 hpf and 27 hpf. We chose the 21 hpf time point because this is when EJC mutant embryos begin to show visible phenotypes and reduced Rbm8a and Magoh protein levels (Fig. 2C and 2D) but do not yet display extensive cell death. We chose the 27 hpf time point because this is when EJC mutant embryos have reliably low Rbm8a and Magoh protein levels as well as motor axon growth defects. However, because *magoh* mutant embryos display extensive necrosis at 27 hpf (Fig. S2B), we only focused on RNA-Seq from the less necrotic *rbm8a* mutant embryos at this time point. We generated three biological replicates of ribosomal RNA-subtracted total RNA-Seq datasets from each mutant (*rbm8a* mutant at 21 hpf and 27 hpf, and *magoh* mutant at 21 hpf, Table S1) and their wild-type siblings (a mixture of wild-type and heterozygous embryos). A differential gene expression analysis using DESeq2 identified gene-level expression changes in the two mutant embryos compared to their wild-type siblings. As expected, *rbm8a* and *magoh* transcripts are downregulated in the respective mutant embryos (Fig. 4A-C). We compared genes that are significantly altered (fold-change > 1.5, false discovery rate (FDR) < 0.05) among the different mutant embryos and time points. When 21 hpf *rbm8a* and *magoh* mutant embryos are compared, a significant number of genes are commonly up- or down-regulated (Fig. 4D, 103 upregulated and 29 downregulated). Similarly, a significant number of genes are commonly upregulated between *rbm8a* mutant embryos at 21 and 27 hpf (Fig. 4E). The modest overlap we observe between differentially expressed genes in EJC mutant embryos could be due to differences in developmental timing, due to variable degree of protein depletion, or due to EJC-independent functions of individual proteins. Despite changes in transcript levels in the two mutant embryos, no reproducible splicing changes are identified in these datasets using DEX-Seq, an algorithm that estimates differential exon usage (data not shown).

**Figure 4.**
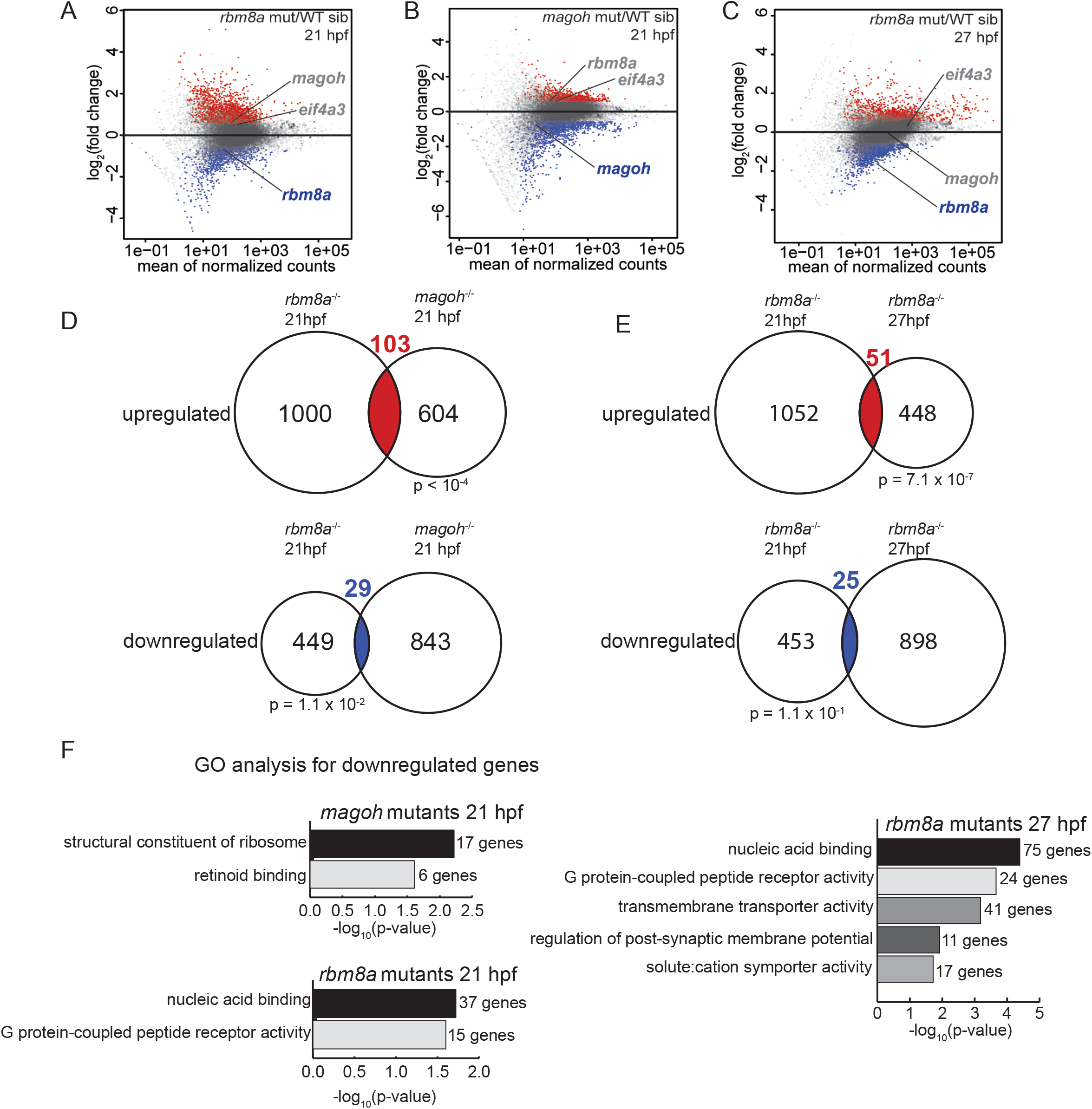
Gene expression changes in *rbm8a* and *magoh* mutant embryos. A-C. MA plots (M: log ratio; A: mean average) showing genes that are upregulated (fold change > 1.5 and FDR < 0.05) (red), downregulated (fold change < 1.5 and FDR < 0.05) (blue), or unchanged (gray) in *rbm8a* mutant embryos compared to WT siblings at 21 hpf (A), *magoh* mutant embryos compared to WT siblings at 21 hpf (B), and *rbm8a* mutant embryos compared to WT siblings at 27 hpf (C). *rbm8a*, *magoh*, and *eif4a3* are labeled in each plot with label colors signifying no change (gray) or downregulation (blue). D. Venn diagrams showing the overlap between genes that are upregulated (top) and downregulated (bottom) in *rbm8a* mutant embryos at 21 hpf (left) and *magoh* mutant embryos at 21 hpf (right). Hypergeometric test p-values are below each comparison. E. Venn diagrams as in (D) comparing upregulated and downregulated genes in *rbm8a* at 21 (left) and 27 hpf (right). F. PANTHER14.0 (70) gene ontology (GO) term overrepresentation analysis of genes downregulated in *rbm8a* and *magoh* mutant embryos at indicated times. All significant terms (Benjamini-Hochberg corrected p-value < 0.05) are shown for each set. The number of genes in each term is indicated at the right of each bar.

We next determined if genes with shared functions are enriched among differentially-expressed genes in EJC mutant embryos. Except for a handful of cell death regulators and effectors, none of the upregulated genes in either EJC mutant identify a functionally-related class of genes, suggesting that the proteins they encode perform a variety of functions. The upregulated cell death genes include *tp53*, *tp53-inp1*, and *casp8* (the latter upregulated only in *rbm8a* mutant embryos), which is consistent with the cell death observed in mutant embryos (Fig. 2B and S2). In contrast to upregulated genes, downregulated genes in each EJC mutant are enriched in specific GO terms. In *rbm8a* mutant embryos, downregulated genes at both 21 and 27 hpf are significantly enriched for genes encoding proteins with G-protein coupled receptor (GPCR) or nucleic acid binding activities (Fig. 4F). In *magoh* mutant embryos at 21 hpf, downregulated genes are significantly enriched for the retinoid binding GO term, which includes several GPCRs. Another functionally related group of genes downregulated in *magoh* mutant embryos at 21 hpf are genes encoding structural constituents of the ribosome (Fig. 4F). This latter class is also downregulated in mouse *magoh* heterozygotes (11), highlighting the importance of *magoh* in ribosomal gene expression.

### *rbm8a* and *magoh* mutant embryos have defects in NMD

Because translation termination upstream of exon-exon junctions was previously shown to trigger NMD in zebrafish embryos (37), we expected that one group of upregulated transcripts in EJC mutant embryos would be mRNAs containing premature termination codons (PTC) or “natural” NMD targets containing a 3’ UTR intron (3′UI) or an upstream open reading frame (uORF). To identify whether NMD targets are enriched among upregulated genes in EJC mutant embryos, we first compared genes upregulated in EJC mutant embryos to those upregulated in zebrafish *upf1* morphants at 24 hpf (38). A significant number of genes are shared between 24 hpf *upf1* morphants and *magoh* mutant embryos at 21 hpf (39 out of 707), and *rbm8a* mutant embryos at 21 hpf (45 out of 1103, data not shown) and at 27 hpf (44 out of 499) (Fig. S3A). Importantly, NMD targets previously validated in zebrafish (e.g. *isg15*, *atxn1b*, *bbc3*) (38) and mRNAs predicted to undergo NMD (e.g. *upb1,* contains a 3′UI) are among these shared genes. We also generated an independent dataset of Upf1-regulated transcripts from zebrafish morphants at an earlier timepoint (12 hpf) using RNA-Seq (in duplicate, Fig. S3B, Table S1) to avoid secondary targets upregulated due to significant cell death in *upf1* morphants (37). We find that upregulated genes in 12 hpf *upf1* morphants (Fig. S3C) show a modest but significant overlap with upregulated genes in the previously published 24 hpf *upf1* morphant dataset (Fig. S3C); the overlap also includes three of the five previously validated NMD targets (*isg15*, *atxn1b* and *bbc3*) (38). Significant overlap is also observed among upregulated genes in 12 hpf *upf1* morphants and 21 hpf *magoh* mutant embryos (32 out of 707, Fig. 5A, Table S2), 21 hpf *rbm8a* mutant embryos (33 out of 1103, data not shown), and 27 hpf *rbm8a* mutant embryos (65 out of 499, Fig. 5A, Table S2). Because the observed changes in *rbm8a* mutant embryos at 21 hpf were similar to but more modest than in *magoh* and *rbm8a* mutant embryos at 21 hpf and 27 hpf, respectively, we focused all subsequent analyses on the latter two EJC mutant datasets. At least one-third of all genes upregulated >1.5 fold upon *upf1* knockdown (FDR < 0.05) also show a >1.5-fold increase (FDR < 0.05) in either 21 hpf *magoh* or 27 hpf *rbm8a* mutant embryos with 14 genes being significantly upregulated in all three datasets (Fig. 5A, Table S2). Globally, genes upregulated >1.5 fold in EJC mutant embryos (FDR < 0.05), as compared to unchanged genes, also show a positive fold-change in *upf1* morphants at both 12 hpf and 24 hpf (Fig. 5B-C; Fig. S3D-G). This suggests that while only a small set of genes is significantly affected upon depletion of the EJC or Upf1, a much larger shared set of genes show an increase in abundance.

**Figure 5.**
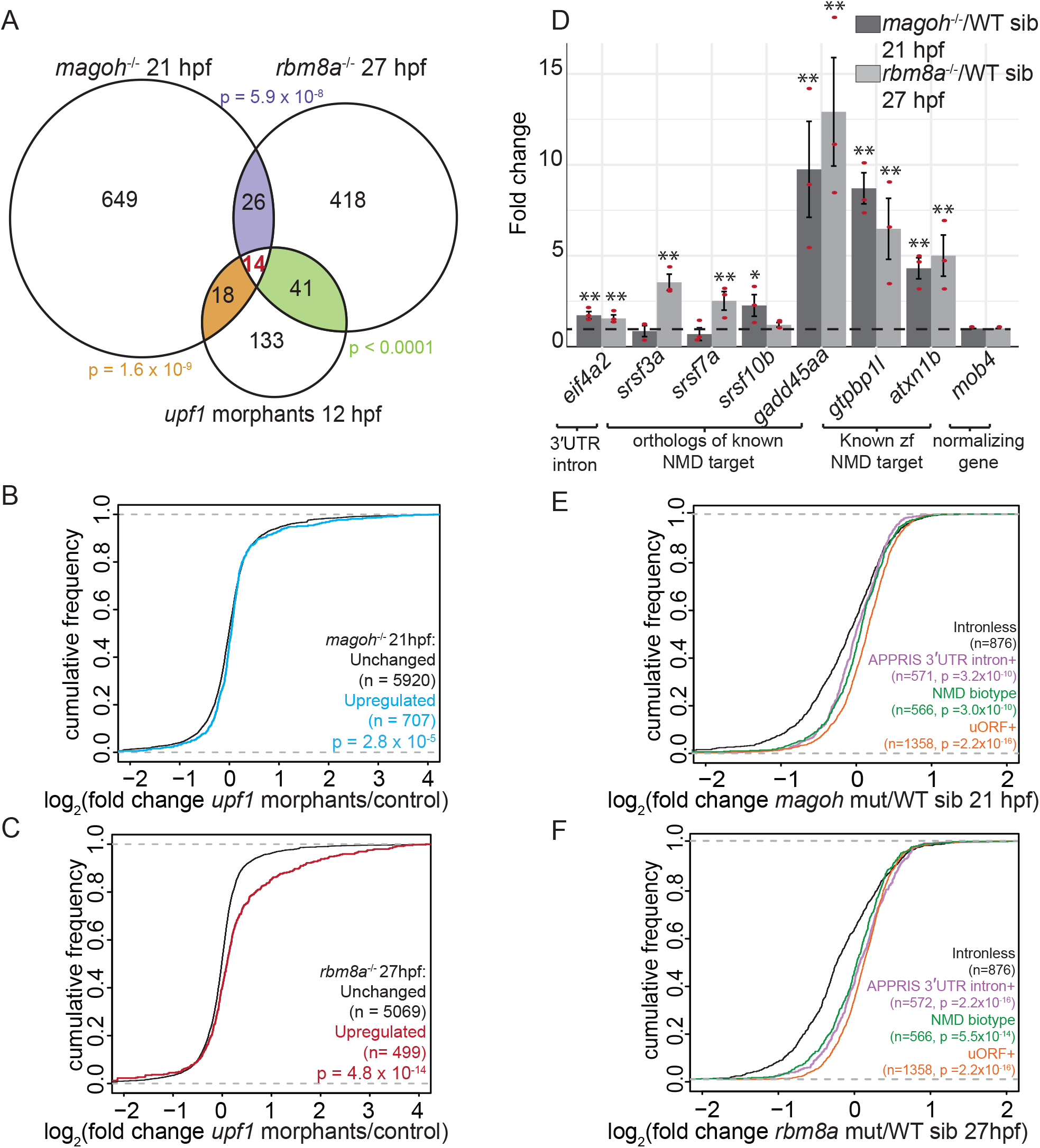
Genes upregulated in EJC mutant embryos are also regulated by Upf1 and contain NMD-inducing features. A. Venn diagram showing the overlap of significantly upregulated genes in EJC mutant embryos and *upf1* morphants. Each overlap and its corresponding hypergeometric test-based p-value are color-coded. B. Cumulative distribution frequency (CDF) plot showing the fold changes in *upf1* morphants (12 hpf) of genes upregulated in *magoh* mutant embryos at 21 hpf (blue) compared to unchanged genes (black). Kolmogorov-Smirnov (KS) test p-value for differences in fold changes between the two groups is indicated on the bottom right. C. CDF plot as in B for genes upregulated in *rbm8a* mutant embryos at 27 hpf (red) compared to unchanged genes (black). D. Quantitative RT-PCR (qRT-PCR) analysis showing fold change of select NMD target transcripts (x-axis) compared to control (*mob4*) transcript in *magoh* mutant embryos at 21 hpf compared to wild-type siblings (dark gray bars) and in *rbm8a* mutant embryos at 27 hpf compared to wild-type siblings (light gray bars). The selected genes either contain a 3′UTR intron and/or have orthologs that are known NMD targets or were previously shown to be zebrafish Upf1 targets (38). Red dots: the value of each individual replicate. Error bars: standard error of means. Horizontal black dashed line: fold change = 1. Welch t-test p-values are indicated by asterisks (** p-value < 0.05; * p-value < 0.1). E. CDF plot showing the fold changes in 21 hpf *magoh* mutant embryos for genes that contain 3′UTR introns (APPRIS 3′UTR intron, mauve), uORF (orange), defined in Ensembl as NMD-biotype (green) compared to intronless genes (black). KS test p-value for differences in distribution of fold changes between intronless genes and each of the particular groups is indicated on the bottom right. F. CDF plot as in E showing the fold changes in *rbm8a* mutant embryos at 27 hpf.

To further confirm that predicted EJC-dependent NMD targets are indeed affected in EJC mutant embryos, we quantified relative levels of select transcripts that are orthologous to well-known NMD targets (*srsf3a, srsf7a, srsf10b,* and *gadd45aa*), or are upregulated in *upf1* morphants (e.g. *gtpbp1l*, *atxn1b*). All of these transcripts are robustly upregulated in at least one of the EJC mutant backgrounds compared to wild-type siblings, and some (*eif4a2*, *pik3r3a*, *gadd45aa*, *gtpbp1l*, *atxn1b*) are upregulated in both EJC mutant backgrounds (Fig. 5D). These data further confirm that the EJC is required for Upf1-mediated downregulation of NMD targets in zebrafish embryos.

We next tested if different known classes of NMD targets (PTC-, 3′UI-or uORF-containing mRNAs) are upregulated in EJC mutant embryos. We identified 589 genes that encode transcripts annotated as ‘NMD biotype’ in Ensembl due to the presence of PTCs. When compared to intron-less protein-coding genes, 566 genes encoding NMD biotype transcripts show an increase in abundance in EJC mutant embryos (Fig. 5E-F). We also identified 582 genes that encode transcript isoforms with 3′UIs > 50 nts from stop codons and are reliably detectable according to the APPRIS database (39). Similar to the NMD biotype group, the genes encoding APPRIS-supported 3′UI-containing transcripts are also increased in abundance in EJC mutant embryos (Fig. 5E-F). Another feature known to induce EJC-dependent NMD is uORFs. Based on a published ribosome footprinting dataset (40, 41) we identified 1525 genes encoding transcripts with ribosome-occupied uORFs. When compared with a control set of intron-less protein-coding genes, genes containing functional uORFs show a significant positive fold change in EJC mutant embryos (Fig. 5E-F). Overall, we conclude that the ability of PTCs, 3′UIs and uORFs to trigger EJC-dependent NMD is conserved in zebrafish.

### Transcripts with stop codon-proximal 3′UTR introns are upregulated upon loss of EJC and Upf1 function

Current models of vertebrate NMD state that translation termination must occur at least 50 nts upstream of the last exon-exon junction for an mRNA to undergo EJC-dependent NMD. To date, a T-cell receptor *β* (*TCR-β*) derived transcript is the only known exception to this “50-nt rule” (42). Surprisingly, we noticed that among the 14 transcripts that are commonly upregulated in *rbm8a* mutant embryos, *magoh* mutant embryos, and *upf1* morphants (Fig. 5A), three (*foxo3b*, *phlda3* and *nupr1a*) are encoded by genes that contain a 3′UI where the distance between the stop codon and the intron is less than 50 nts. For *foxo3b* and *nupr1a*, the human orthologs also contain a 3′UI < 50 nts downstream of the stop codon. This observation raises an intriguing possibility that additional mRNAs exist that defy the 50-nt rule, and that a proximal 3′UI (< 50 nts distance between intron and upstream stop codon; Fig. 6A) may represent a previously unrecognized NMD-inducing feature. Using Ensembl GRCz10 transcript annotations, we identified 861 zebrafish genes that encode transcripts with proximal 3′UIs and 582 genes that encode transcripts with distal 3′UIs (3′UIs ≥ 50 nts downstream of the stop codon; Fig. 6A) (Fig. S4A, Table S3). Interestingly, proximal 3′UI-containing genes encode proteins which are enriched for mRNA binding and mRNA splicing factor functions (Fig. S4B), two functional groups that are well-recognized to be regulated by EJC-dependent NMD (43, 44). We find that 3.5-8% (70/854 in *rbm8a* 27 hpf, 60/854 in *magoh* 21 hpf and 21/597 in *upf1* morphants 12 hpf) of all proximal 3′UI-containing genes, detectable in our datasets, are ≥ 1.5 fold upregulated in all three EJC mutant embryos (Fig. 6B, 6C and S4C) and in *upf1* morphants (Fig. S4D). To further confirm that proximal 3′UI-containing genes are indeed targets of NMD, we focused on a subset (*foxo3b*, *cdkn1ba*, and *phlda2*) that are upregulated in *upf1* morphants and at least one EJC mutant, and where the existence of a proximal 3′UI is conserved in several other vertebrates including humans. After treating embryos with the NMD inhibitor NMDI14 (45), we found that *foxo3b*, *cdkn1ba*, and *phlda2* transcripts are 2-to-8 fold upregulated, just like *eif4a2,* a distal 3′UI-containing transcript, and *atxn1b*, a previously validated (38) NMD target (Fig. 6D). Thus, a subset of genes with a 3′UI in a stop codon-proximal position appear to defy the 50-nt rule as their encoded mRNAs are targeted for EJC-dependent NMD. Interestingly, we did not observe any correlation between the distance of the 3′UI from the stop codon and the degree of fold change observed in EJC mutant or *upf1* morphants (Fig. 6B-C, S4C-D).

**Figure 6.**
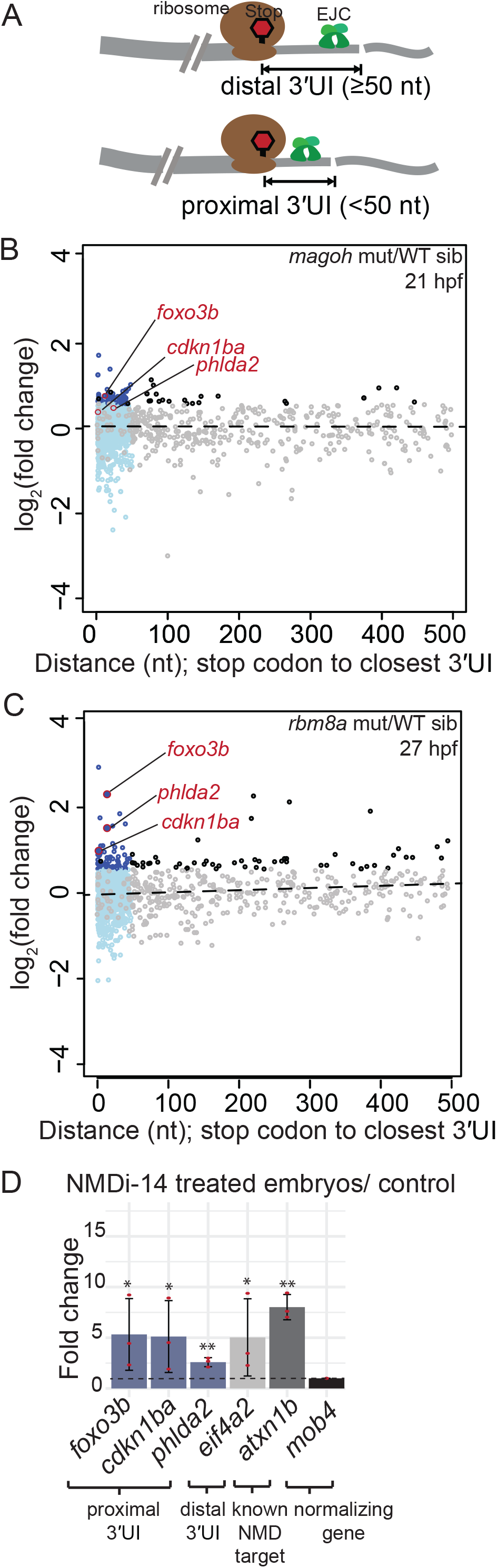
Transcripts encoded by genes with a proximal 3′UTR intron are upregulated in EJC mutant and in NMDI14-treated embryos. A. Top: Schematic illustrating genes with 3′UTR introns (3′UI) where the distance between the stop codon and the 3′UI is equal to or greater than 50 nts. Such 3′UI are classified as distal. Bottom: Schematic illustrating genes with 3′UI where the distance between the stop codon and 3′UI is less than 50 nts. Such 3′UI are classified as proximal. The ribosome, stop codon and EJC are labeled in the top panel. B. A scatter plot showing fold change (FC) for genes with proximal 3′UI (dark blue: FC > 1.5 and light blue: FC < 1.5) and distal 3′UI (black: FC > 1.5 and gray: FC < 1.5) in *magoh* mutant embryos at 21 hpf compared to wild-type siblings. Genes encircled in red also contain a proximal 3′UI in mouse and human, and were independently validated in (D). Gray and black dots in the < 50 nts region represent genes with one proximal and one or more distal intron. C. A scatter plot as in B showing fold changes for *rbm8a* mutant embryos at 27 hpf compared to wild-type siblings. D. qRT-PCR analysis showing fold changes for proximal 3′UI-containing genes (blue bars), a distal 3′UI-containing gene (light gray bar), and a validated zebrafish Upf1-regulated gene (dark gray bar) compared to the control gene (black bar) in zebrafish embryos treated with NMDI14 from 3-24 hpf. Red dots: the value of each individual replicate. Error bars: standard error of means. Horizontal black dotted line: fold change = 1. Welch t-test p-values (** p-value < 0.05; * p-value < 0.1).

### Loss of function of *foxo3b*, a proximal 3′UI-containing gene upregulated in EJC mutant embryos, partially rescues motor axon outgrowth

The zebrafish *foxo3b* gene, whose vertebrate orthologs also contain a proximally-located 3′UI (Fig. 7A), encodes a transcript that is significantly upregulated in both EJC mutant and *upf1* morphants (Fig. 6B-C, S4C-D, and S5A). Conservation of proximal intron position suggests that the stop codon-proximal 3′UI may be important for regulation of *foxo3b* expression. Like *foxo3b* transcript, we find that Foxo3b protein is upregulated 2.8-fold in 21 hpf *magoh* mutant embryos (Fig. 7B) and 1.3-fold in 27 hpf *rbm8a* mutant embryos (Fig. S5B) as compared to wild-type siblings. Furthermore, five known Foxo3b transcriptional target genes (46) are also upregulated in *magoh* and *rbm8a* mutant embryos (Fig. S5C). To test if Foxo3b upregulation contributes to EJC mutant phenotypes, we obtained a previously described *foxo3b* null allele, *foxo3b^ihb404^* (47, 48), generated *magoh; foxo3b* and *rbm8a; foxo3b* doubly heterozygous adults, and examined muscle and motor neuron development in single, double, and compound mutant embryos. As expected, motor axon outgrowth in *foxo3b* mutant embryos is indistinguishable from wild-type siblings (data not shown); in contrast, as noted above, motor axons barely extend beyond the horizontal myoseptum in EJC mutant embryos (Fig. 3E-G, 7G-H, and S5H-I). Strikingly, we find that heterozygous and homozygous loss of *foxo3b* in EJC mutant embryos leads to significantly longer motor axons that extend well beyond the horizontal myoseptum (Fig. 7I-K and S5J-L), but not as far as in wild-type embryos. Despite significant rescue of motor axon outgrowth, neuromuscular junction formation (Fig. 7G-J and S5H-K) and myofiber organization (Fig. 7C-F and S5D-G) are not restored in *magoh; foxo3b* or *rbm8a; foxo3b* double mutant embryos. Thus, we hypothesize that Foxo3b repression via EJC-dependent NMD is important for motor axon outgrowth, but that regulation of other targets is required for proper muscle development.

**Figure 7.**
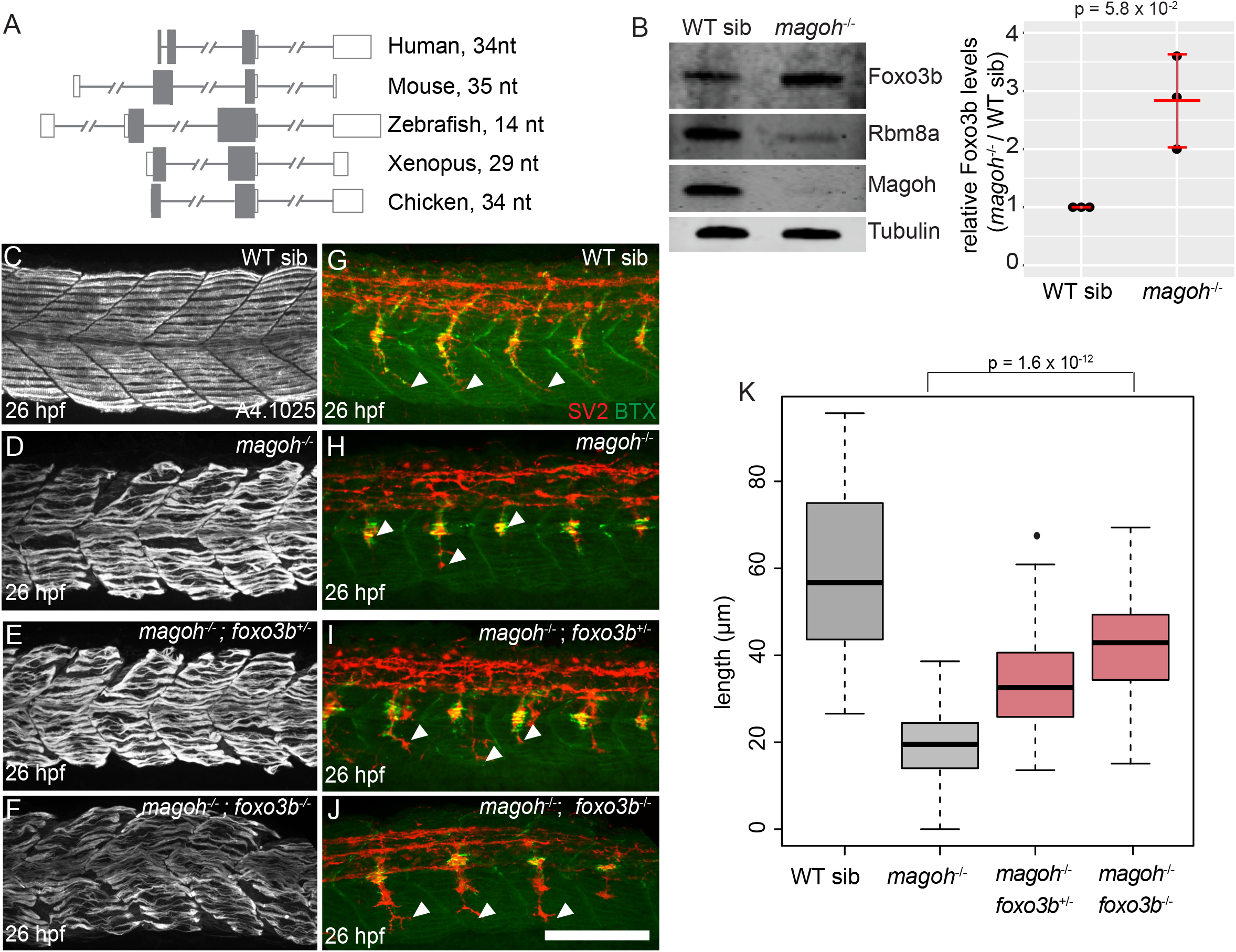
Partial or complete loss of *foxo3b* in *magoh* mutant embryos rescues motor neuron outgrowth defects. A. Illustration showing *foxo3b* gene structure in multiple vertebrates. The distance between the stop codon and the proximal 3′UTR intron is on the right. Open rectangles: UTRs, filled rectangles: coding region, gray lines: introns (hash marks denote shortened intron sequences). B. Left: Western blot showing protein levels in wild-type sibling (left) and *magoh* mutant (right) embryos at 21 hpf. Right: a dot plot showing Foxo3b levels normalized to tubulin levels in *magoh* mutant embryos and WT siblings at 21 hpf in three biological replicates. (N= 5 embryos per genotype per replicate). Error bars: standard error of means. C-F. Confocal images showing Myh1 immunofluorescence using anti-A4.1025 in somites 12-16 of WT sibling (C), and *magoh*^-/-^ mutant (D), *magoh*^-/-^; *foxo3b*^+/-^ mutant (E), and *magoh*^-/-^; *foxo3b*^-/-^ mutant (F) embryos. (N = 13 embryos/genotype). Scalebar in J (for panels C-J) is 100 nm. G-J. Merged confocal images showing motor neurons (red; detected by anti-SV2 staining) and acetylcholine receptors (green; detected by alpha-bungarotoxin staining) in somites 12-16 of WT sibling (G), *magoh*^-/-^ mutant (H), *magoh*^-/-^; *foxo3b*^+/-^ mutant (I), and *magoh*^-/-^; *foxo3b*^-/-^ mutant (J) embryos. Neuromuscular junctions in the merged image are yellow. White arrowheads point to the distal end of the motor neuron. (N = 13 embryos per genotype). Scalebar in J (for panels C-J) is 100 nm. K. Boxplots showing quantification of motor neuron length in embryos of genotypes indicated along the x-axis (4 motor neurons/embryo and 13 embryos/genotype).

### Proximal 3′UI-containing genes are regulated by NMD in human and mouse cells

We surveyed human and mouse genomes for prevalence and conservation of proximal 3′UIs. Like in zebrafish, proximal 3′UI-containing genes outnumber distal 3′UI-containing genes in human (Fig. S6A and Table S3, 1239 proximal 3′UI genes, 489 distal 3′UI genes) and mouse (Fig. S6B and Table S3, 921 proximal 3′UI genes, 649 distal 3′UI genes). Except for over-representation immediately downstream of the stop codon, 3′UI position appears randomly distributed within the stop-codon proximal window, and 3′UI occurrence precipitously drops after the 50 nts position (Fig. S4A, S6A and S6B). As in zebrafish, human and mouse proximal 3′UI-containing genes are enriched in mRNA binding function (Fig. S6C, D). A cross-comparison of zebrafish, mouse, and human proximal 3′UI-containing genes identified 167 genes where the proximal position of 3′UI is conserved in all three organisms suggesting that proximal 3′UIs could be important regulatory elements. These genes show a significant interaction network amongst themselves (Fig. 8A, p-value = 0.02), and are enriched for genes encoding proteins with RNA recognition motifs and with roles in neural development and disease (Fig. 8B).

**Figure 8.**
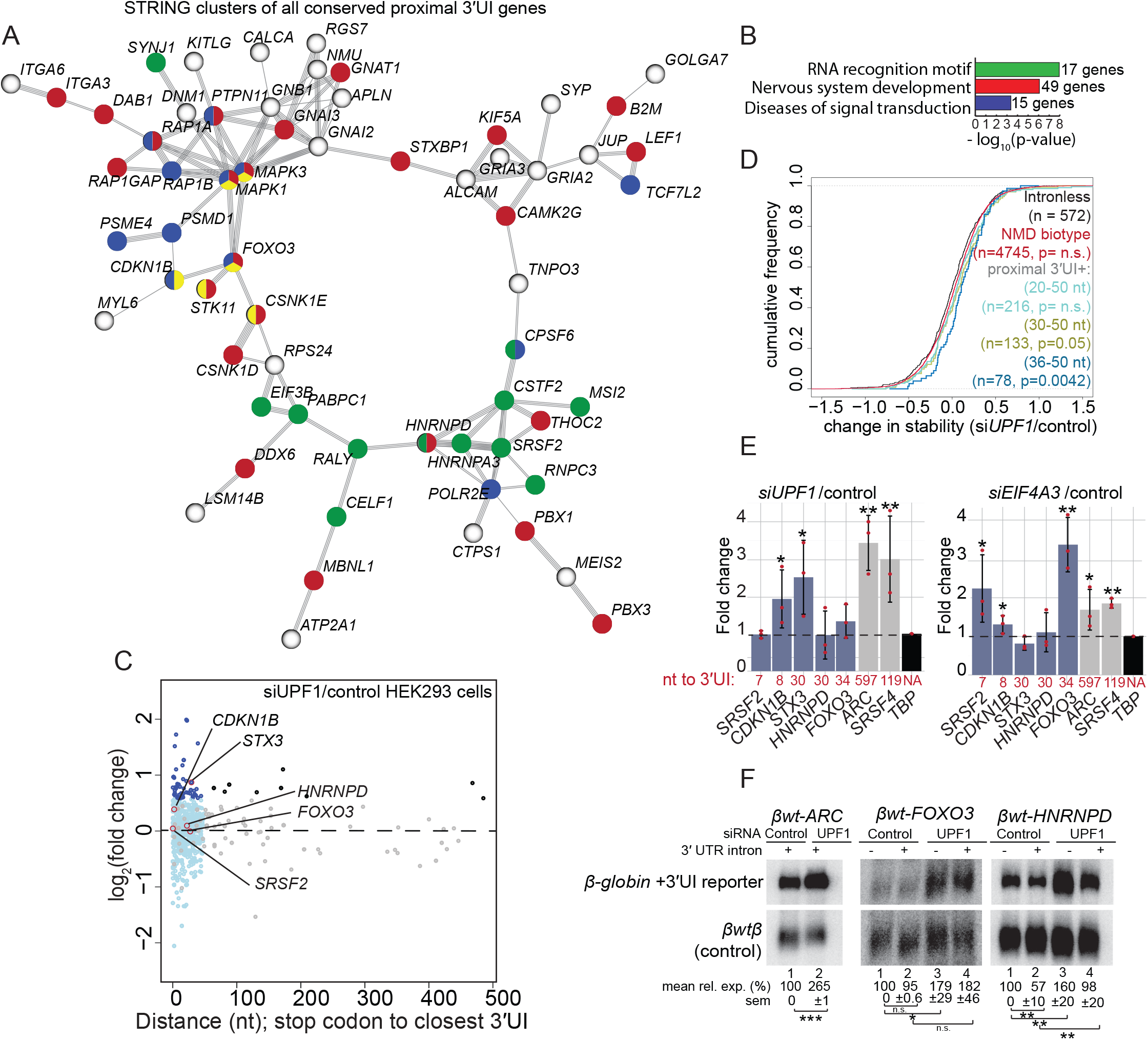
Proximal position of 3′UTR introns is conserved in many vertebrate genes and such introns can induce NMD in human cells. A. A major interaction cluster predicted by STRING network analysis of genes with a shared proximal 3′UI in zebrafish, mouse and human. Nodes are colored by gene/protein function: nervous system (red), presence of RNA recognition motif (green), diseases of signal transduction (blue), FoxO signaling pathway (yellow). (167 nodes and 127 edges in total, PPI enrichment p-value = 0.02). B. Gene ontology enrichment analysis of all 167 genes with conserved 3’UI proximal positioning. The most significant GO term within the following functional categories has been shown: Interpro domains, Biological process and Reactome pathways. C. A scatter plot showing fold changes for all proximal 3′UI genes (dark blue: FC > 1.5 and light blue: FC < 1.5) and all distal 3′UI genes (black: FC > 1.5 and gray: FC < 1.5) in *UPF1* knockdown HEK293 cells compared to control cells using previously published data (49). Genes encircled in red are further tested in (E). Gray and black dots in the < 50nts region represent genes with one proximal and one or more distal intron. D. CDF plot showing change in mRNA stability for different classes of NMD targets and intron-less genes upon *UPF1* knockdown in HEK293 cells (data derived from 46). The gene classes are as follows: proximal 3′UI-containing genes where distance is 20-50 nts (sky blue), 30-50 nts (olive green) and 36-50 nts (dark blue), Ensembl-annotated NMD-biotype genes (red) and intron-less genes (black). KS test p-value for comparison of NMD targets to intron-less genes is indicated in the same color. E. qRT-PCR analysis showing fold changes for proximal 3′UI-containing genes (*FOXO3*, *CDKN1B*, *HNRNPD, STX3* and *SRSF2*) and distal 3′UI-containing genes (ARC and *SRSF4*) upon *UPF1* (left) and *EIF4A3* (right) knockdown in HCT116 cells. The distance between stop codon and 3′UI for every 3′UI-containing gene is indicated below each bar. *TBP* is the normalizing gene used for qPCR analysis. Welch t-test p-values are indicated using asterisks (** p-value < 0.05 and * p-value < 0.1). F. Northern blots showing expression levels of *β-globin* reporters upon control or *UPF1* knockdown in HeLa Tet-off cells. Indicated on top are reporter names, siRNA transfection condition, and whether or not a 3′UTR intron is present in the reporter. Co-transfected wild-type *β-globin* (*βwtβ*) mRNA serves as normalizing control within each lane. Relative expression (% of lane 1 for each reporter) and standard error (based on three replicates) are indicated below each lane.

To test if human and mouse proximal 3′UI-containing genes are also regulated by NMD, we analyzed publicly available RNA-seq datasets of human and mouse cell lines depleted of key NMD factors. We find that a subset of proximal and distal 3′UI-containing genes are similarly upregulated in *UPF1*-depleted HEK293 cells (Fig. 8C) (49) and human ESCs (Fig. S6E) (50), and in *Smg6^-/-^* knockout mouse ESCs (Fig. S6F) (51). In addition to steady state RNA levels, analysis of RNA stability in HEK293 cells (49) showed that UPF1 knockdown progressively increases transcript stability of proximal 3′UI-containing genes grouped based on increasing distance from the stop codon (20-50 nts, 30-50 nts, and 36-50 nts) (Fig. 8D). Notably, the proximal 3′UI-containing genes where the intron is ≥ 36 nts from the stop codon are the most significantly stabilized. To further confirm UPF1 and EJC dependence of proximal 3′UI NMD targets in human cells, we knocked down *UPF1* and *EIF4A3* in a human colorectal carcinoma cell line (HCT116), and assessed levels of transcripts that are upregulated in at least one of the human/mouse datasets analyzed above (*SRSF2, CDKN1B, STX3, HNRNPD* and *FOXO3*) and where proximal position of the 3′UI is conserved (with the exception of *STX3*; Fig. 8A). We find that *CDKN1B* is upregulated upon *EIF4A3* and *UPF1* knockdown like transcripts encoded by distal 3′UI-containing genes (*ARC and SRSF4)* (Fig. 8E). *SRSF2, STX3* and *FOXO3* are upregulated either upon *UPF1* or *EIF4A3* knockdown but not under both conditions. *HNRNPD*, on the other hand, remains unchanged in HCT116 cells upon *UPF1* or *EIF4A3* knockdown (Fig. 8E). Thus, proximal 3′UI-containing transcripts are regulated by EJCs and UPF1 but exhibit differential sensitivities to loss of EJC-dependent NMD components possibly due to transcript and/or cell-type specific differences in NMD.

To test if proximal 3′UIs are sufficient to induce EJC-dependent NMD, we generated human *β-globin* mRNA-based reporters by replacing the *β-globin* 3′UTR with proximal 3′UI-containing 3′UTRs from human *FOXO3* and *HNRNPD* genes. In the case of *FOXO3*, only the first ∼0.9 kb of the ∼2.7 kb 3′UTR was used to avoid triggering NMD due to 3′UTR length (19). We also created matching control reporters lacking the 3′UTR intron. As a positive control for 3′UI-dependent NMD, we inserted the *ARC* 3′UTR (*βwt-ARC*), which contains two distal NMD-inducing 3′UIs (52). All reporters show expected RNA processing patterns in HeLa cells including 3′UI splicing (data not shown). If a proximal 3′UI promotes EJC-dependent NMD, 3′UI-containing reporter transcripts are expected to be less abundant compared to the 3′UI-lacking reporters under normal conditions, and to be upregulated upon *UPF1* knockdown. Indeed, we observe such a trend for *βwt-HNRNPD* in HeLa cells (Fig. 8F and S4H, compare lanes 1 and 2 in *βwt-HNRNPD* panel). Surprisingly, both *HNRNPD* and *FOXO3* 3′UTRs are able to induce UPF1-dependent downregulation even in the absence of a proximal 3′UI. However, unlike *HNRNPD* 3′UI, the *FOXO3* 3′UI cannot further downregulate the *β-globin* reporter (Fig. 8F). Overall, we conclude that, contrary to the prevailing dogma, certain proximal 3′UIs are capable of inducing NMD.

## Discussion

Our work sheds light on EJC assembly and function during early zebrafish development, highlights the conserved role of the EJC in NMD among vertebrates, and demonstrates that a proximal 3′UI can function as an NMD-inducing feature that modulates gene expression.

### Conserved features of EJC assembly in vertebrates

To our knowledge, our work is the first to illuminate EJC occupancy in a whole vertebrate organism, and reveals conserved features of EJC assembly. In zebrafish embryos, the highly conserved EJC core proteins show predominant binding ∼24 nts upstream of exon-exon junctions, with exon to exon variability (Fig. 1), as observed in human cells (29, 30). In addition to the canonical position, the zebrafish EJC is also detected at non-canonical positions (Fig. 1), just like in human cells (29–31,33). Interestingly, EJC binding to non-canonical regions appears to be unique to vertebrates as the *Drosophila* EJC binds almost exclusively at canonical positions (32). The precise molecular basis of non-canonical EJC occupancy, and its impact on post-transcriptional gene regulation remain important outstanding questions for the future.

### Loss of the EJC causes tissue-specific defects and embryonic lethality in zebrafish

In either of the *rbm8a* and *magoh* zebrafish mutant embryos, we observe a strong co-depletion of both Rbm8a and Magoh proteins (Fig. 2C and 2D). This simultaneous deficiency of *rbm8a* and *magoh* function in zebrafish embryos likely impairs EJC function leading to rapid emergence of developmental defects, which progressively worsen and eventually cause embryonic death by 2 dpf. The earliest phenotype observed in EJC mutant embryos is neural cell death (observed by acridine orange staining) in the brain and spinal cord at 19 hpf (Fig. S2A); by 21 hpf, brain necrosis is also morphologically apparent (Fig. S2B). The neural cell death phenotype is similar to that seen in mouse EJC heterozygous mutant embryos (10, 11), and is consistent with the microcephaly phenotype in human patients heterozygous for hypomorphic *RBM8A* and *EIF4A3* mutations (8, 9). Thus, EJC function during neural development appears to be conserved across vertebrates. The emergence of defects in EJC mutant embryos in discrete lineages such as neural and muscle cells may result from tissue-specific differences in EJC protein functions, activity of EJC regulators, or decay rates of maternally-provided EJC transcript/protein. Future investigation into these possibilities in zebrafish embryos can help explain the tissue-specific nature of EJC-linked defects observed in human syndromes (8, 9). Notably, unlike haploinsufficiency of EJC core components in mouse and human (8–11), heterozygous loss of *rbm8a* or *magoh* in zebrafish does not result in any apparent phenotype indicating a difference in EJC dosage threshold in zebrafish as compared to mammals.

### EJC is a critical component of NMD in zebrafish

Gene expression profiles of *rbm8a* and *magoh* mutant embryos provide insight into molecular functions of the EJC in developing zebrafish embryos. Our finding that a significant fraction of genes upregulated in EJC mutant embryos are also upregulated in *upf1* knockdown embryos (Fig. 5) shows that EJC-dependent NMD is compromised in both *rbm8a* and *magoh* mutant embryos. The overlap between upregulated genes in EJC mutant and *upf1* morphants is small (Fig. 5A and S3A), likely due to differences in developmental timing or due to NMD-independent functions of the proteins. However, the genes within these overlaps include previously validated zebrafish NMD targets and orthologs of known NMD targets in mammals (Fig. 5D and Table S2). Furthermore, several known classes of NMD targets such as PTC-, uORF-, and 3′UI-containing transcripts are significantly upregulated in *rbm8a* and *magoh* mutant embryos (Fig. 5E-F). These findings show that the EJC is critical for both the quality control function (i.e. suppression of aberrant PTC-containing transcripts) and the gene regulatory activity of the zebrafish NMD pathway. These findings highlight the potential of EJC-dependent NMD mechanisms for development/tissue-specific gene regulation, as has also been reported previously (23, 24). An important goal for future work will be to understand how EJC-dependent NMD regulates specific genes to control development. It also remains to be seen if any of the other EJC activities (e.g. mRNA export, RNA localization, translation) also contribute to zebrafish development.

### Proximal 3′UTR introns are a novel NMD-inducing feature

Our surprising discovery that proximal 3′UIs represent a bona fide NMD-inducing signal is supported by multiple lines of evidence. Less than 10 % of all detectable zebrafish proximal 3′UI-containing genes, like distal 3’UI-containing genes, are ≥1.5 fold upregulated in EJC mutant and *upf1* morphant datasets (Fig. 6B, 6C, S4D), and a subset of these are upregulated in zebrafish embryos treated with the NMD inhibitor NMDI14 (Fig. 6D). Further, a subset of proximal 3′UI-containing genes is also upregulated in mouse and human NMD- and EJC-compromised cells (Fig. 8 and S6). Next, the proximal 3′UI in *HNRNPD* 3′UTR can enhance UPF1-dependent downregulation of a *β-globin* reporter RNA (Fig. 8F). The majority of proximal 3′UI-containing genes conserved in human, mouse and zebrafish encode RNA-binding proteins (e.g. *HNRNPD*, *MBNL*), proteins with neuronal functions (e.g. *KIF5A*, *STXBP1*), or both (e.g. *CELF1*, *MSI2*); genes in these classes are well-known for their regulation via 3′UI and NMD (23,43,53). Thus, our findings suggest that among 3′UI-containing genes numerous exceptions exist to the prevailing 50-nt NMD rule.

Apparently, proximal 3′UI genes are variably susceptible to reduced EJC/NMD function. For example, zebrafish *foxo3b* is significantly upregulated in EJC mutant and *upf1* morphants whereas human *FOXO3* is only mildly sensitive to reduced UPF1 levels in cell lines (Fig. 6, S4, 8 and S6) but shows robust upregulation upon EIF4A3 knockdown in HCT116 cells (Fig. 8E). Furthermore, many vertebrate genes with proximal 3′UIs are not upregulated, or are even downregulated, upon diminished EJC/UPF1 function (Fig. 6 and 8). Notably, variable susceptibility to EJC/NMD-deficiency is also seen among distal 3′UI-containing genes. The variability and/or non-responsiveness of some 3′UI-containing genes to EJC/NMD manipulations could be due to detection issues (e.g. low expression at developmental stage/cell type investigated) or due to their variable sensitivity to EJC/NMD protein levels. Downregulation of proximal 3’UI-containing genes could result from indirect effects of compromised EJC/NMD function. It is also possible that some 3′UI-containing genes actively evade NMD. Such a mechanism(s) could operate via 3′UTR-bound proteins as observed for mRNAs with unusually long 3′UTRs (22, 54). This idea is supported by the recent report that HNRNPL binding to 3’UTRs near stop codons can protect mRNAs with long 3′UTRs or downstream EJCs from NMD (55). Such NMD evasion mechanisms may function in cell-type or tissue-specific manner to allow precise RNA regulation in particular cells, tissues or developmental stages.

### How can splicing of a proximal 3′UI lead to NMD?

The prevalent 50-nt rule is presumed to account for the minimum distance required to accommodate a terminated ribosome so that it does not interfere with the downstream EJC. Based on estimates that ribosome footprints at stop codons extend about 9 nts into the 3′UTR (40) and that the EJC 5′ boundary lies about 27 nts upstream of the exon-exon junction (Fig. 1C), a 3′UI located at least 36 nts downstream of a stop codon could induce EJC-dependent NMD via the commonly accepted mechanism. How introns within the first 35 nts of the 3′UTR induce NMD is more perplexing. One possibility is that introns within 35 nts of the stop may trigger NMD via non-canonical EJCs present downstream in the 3′UTR (29, 30). Additionally, EJC-interacting factors such as SR proteins deposited on 3′UTR sequences after 3′UI splicing may also recruit NMD-activating factors (29,30,56,57). Curiously, both proximal and distal 3′UI-containing transcripts show comparable upregulation upon NMD disruption (Figs. 6 and 8). Thus, we speculate that the presence of a 3′UI rather than its 3′UTR position is a stronger determinant of an mRNA’s 3′UTR ribonucleoprotein composition, and hence its NMD susceptibility.

In contrast to 3′UI-mediated NMD, PTC-triggered NMD appears to more strongly adhere to the 50-nt rule (58). Conceivably, some mechanistic differences may exist in induction of NMD at PTCs versus at normal stop codons upstream of a 3′UI. PTC-containing mRNAs are considered aberrant and are targeted for decay to suppress expression of truncated polypeptides whereas 3′UI-containing transcripts are targeted for decay to fine-tune transcript and thus protein levels. We support a hypothesis that translation termination rate at PTCs is likely slow and results in strong NMD induction whereas termination at normal stop codons is rapid and results in weak NMD induction (53).

### EJC-dependent regulation of *foxo3b*, a proximal 3′UI-containing gene, is critical for motor axon outgrowth in zebrafish

Certain genes maintain proximal 3′UIs across vertebrate evolution (Fig. 7 and 8) despite much faster rates of intron loss from 3′UTRs compared to coding region (59, 60), suggesting that proximal 3′UIs play an important role in gene regulation and cellular function. *foxo3b* emerges as one such example that is regulated via EJC-dependent NMD in zebrafish embryos (Fig. 6) and cultured human cells (Fig. 8). This gene encodes a forkhead box transcription factor that acts as a hub for integration of several stress stimuli, and functions in processes such as cell cycle, apoptosis, and autophagy (61, 62). In zebrafish, Foxo3b contributes to survival under hypoxic stress (48), and negatively regulates antiviral responses (47) as well as the canonical *wnt* signaling pathway (63). FOXO3, the mammalian ortholog of Foxo3b, physically interacts with p53, and both act synergistically to induce apoptosis in response to stress (64). In addition, several pro-apoptotic genes (e.g. *bim, bbc3, gadd45a*) are direct FOXO3 transcriptional targets (Fig. S5C) (65, 66). Thus, the regulation of *foxo3b* by EJC-dependent NMD (Fig. 6 and 8) can directly impact cell survival. The elevated levels of Foxo3b in EJC mutant embryos may cause motor neuron outgrowth defects due to dampened Wnt signaling (63, 67), increased neural cell death, or for other cell-autonomous or non-cell-autonomous reasons. Loss of *foxo3b* function in EJC mutant embryos partially reverses these changes (Fig. 7). This finding also correlates well with the rescue of neural apoptosis in mouse EJC mutant embryos upon brain-specific *p53* ablation (10, 11), and reversal of cell death in NMD-defective flies and human cell lines upon reduced activity of *GADD45A* (68, 69). Thus, EJC-mediated regulation of *foxo3b* identifies a new gene in the EJC-NMD-cell survival network and highlights the importance of proximal 3′UIs as post-transcriptional regulatory features during development.

## Supplementary Figure legends

**Figure S1. (related to Figure 1).**
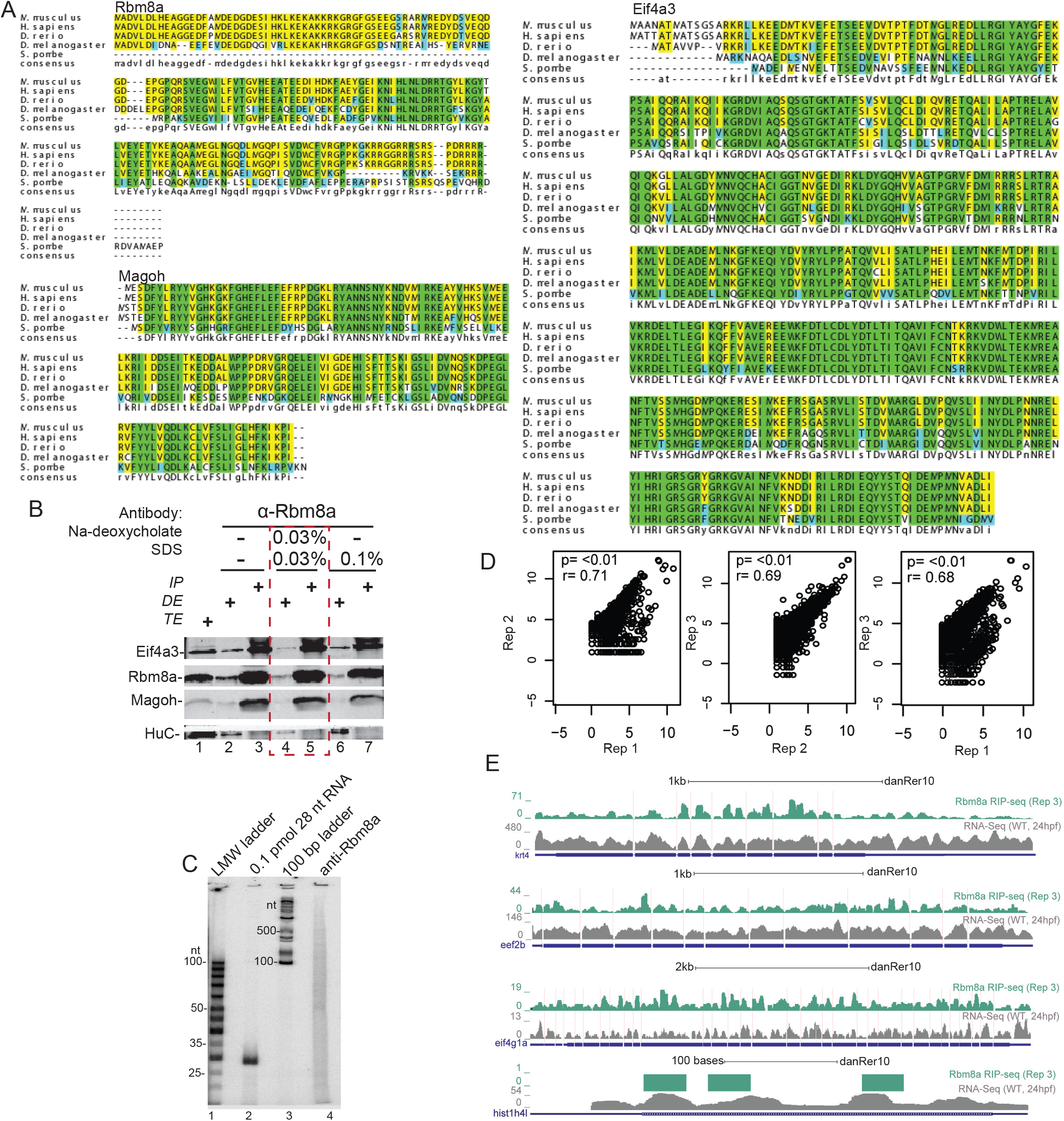
A. Multiple sequence alignments of Eif4a3, Rbm8a and Magoh protein sequences from organisms on the left. Consensus sequence is at the bottom with upper case letters indicating identity and lower case letters indicating similarity. Green indicates complete identity across all species, yellow and blue indicate the identical and unique amino acids in the regions with similarity. Identity between human and zebrafish EJC proteins: Eif4a3 (97%), Rbm8a (93%) and Magoh (100%). B. Western blot detecting proteins listed on the left in RNase I-treated zebrafish embryo total extract (TE, lane 1), depleted extract (DE, lanes 2, 4 and 6) and immunoprecipitates (IP, lanes 3, 5 and 7) with the Rbm8a antibody. Detergents supplemented to increase IP stringency are indicated on top of each lane. Optimized IP condition used in S1C is indicated by the dashed red box. C. Autoradiogram of α^32^P 5′-end labeled RNAs from anti-Rbm8a RIP elution (lane 4) as well as indicated size-markers which include the low-molecular weight single-stranded DNA ladder (lane 1), 0.1 pmol 28 nt synthetic RNA (lane 2) and 100 bp DNA ladder (lane 3). D. Scatter plots comparing read counts for each gene in a pair of RIP-Seq replicates. The replicates (Rep1, Rep2, and Rep3) are indicated on the x- and y-axes. A pseudocount of 0.0001 was added to all genic read counts before log2 transformation. Pearson correlation coefficient and p-value for the correlation test for each comparison is on the top left of each plot. E. Genome browser screenshots showing read coverage of Rbm8a RIP-Seq (only Rep 3, the deepest replicate is shown) in green and RNA-Seq in gray of select highly-expressed genes, *krt4*, *eef2b*, *eif4g1a,* and *hist1h4l (intron-less gene)*.

**Figure S2. (related to Figure 2).**
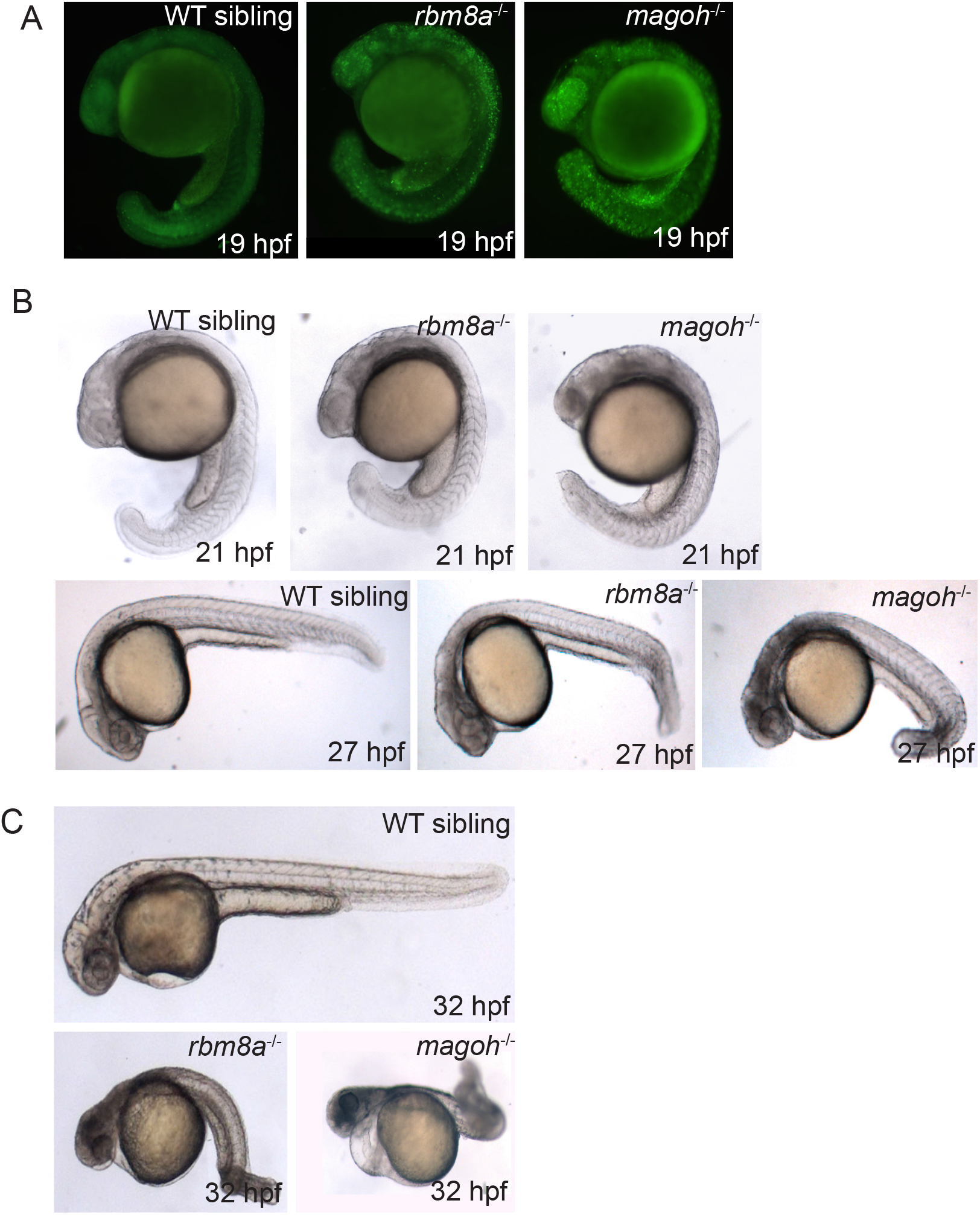
A. Whole mount images of live 19 hpf EJC mutant embryos and WT sibling embryos stained with acridine orange. B. Whole mount images of live EJC mutant embryos and WT siblings at 21 and 27 hpf. C. Whole mount images of live EJC mutant embryos and WT siblings at 32 hpf.

**Figure S3. (related to Figure 5).**
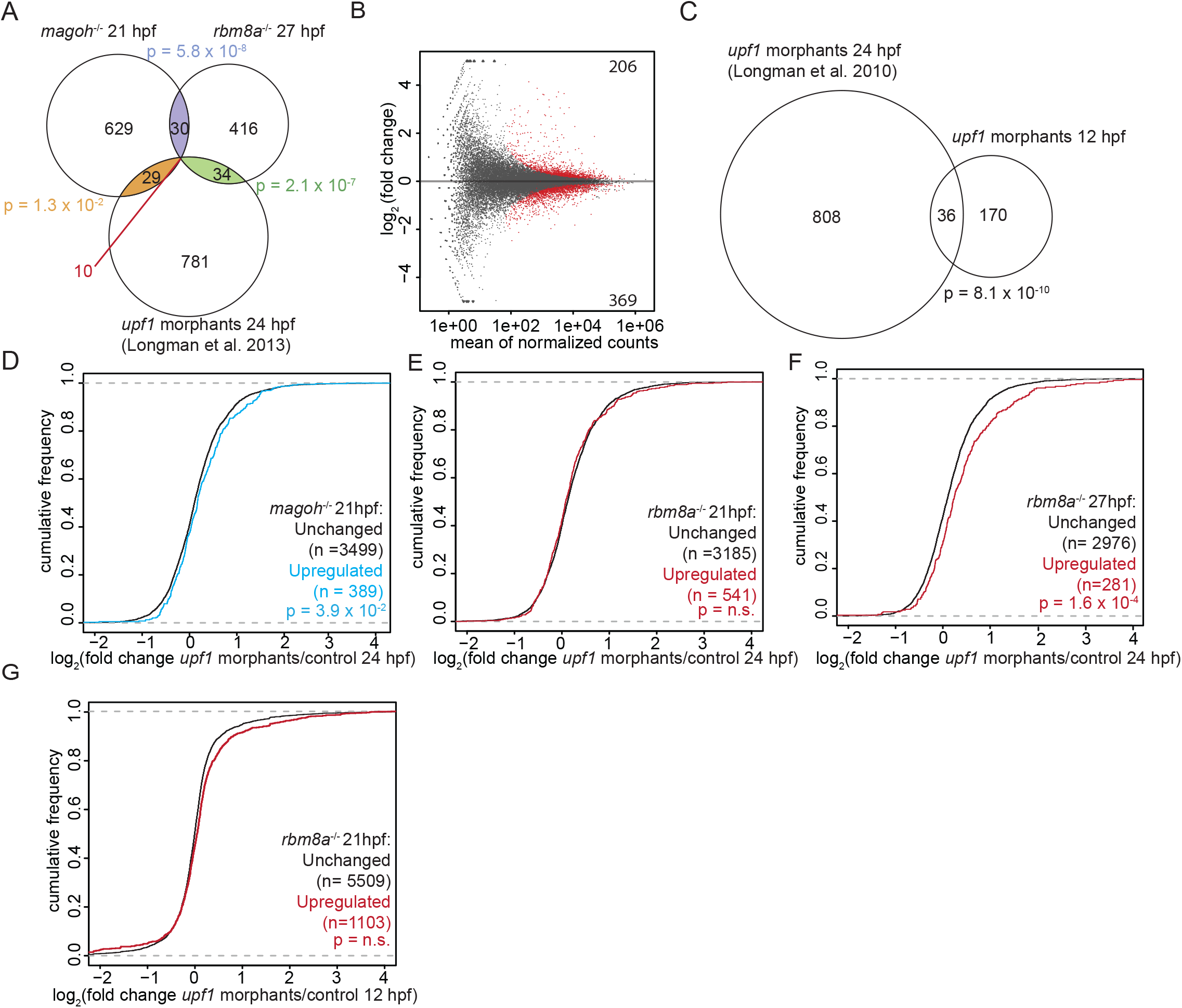
A. Venn diagram showing the overlap of genes that are significantly upregulated in EJC mutant embryos and *upf1* morphant embryos at 24 hpf (38). Hypergeometric test p-values for each comparison are also shown. B. MA plot showing the genes that are altered in expression (fold change > 1.5 and FDR < 0.05; red and unchanged genes in gray) in *upf1* morphant embryos compared to control embryos at 12 hpf. The number of significantly upregulated genes is at the top right and the number of downregulated genes is at the bottom right. C. Venn diagram showing the overlap of significantly upregulated genes in *upf1* morphant embryos at 24 hpf and *upf1* morphant embryos at 12 hpf (38). Hypergeometric test p-value for the comparison is indicated. D. Cumulative distribution frequency (CDF) plot showing the fold changes in *upf1* morphant embryos (24 hpf) (38) of genes upregulated in 21 hpf *magoh* mutant embryos (blue) compared to unchanged genes (black). Kolmogorov-Smirnov (KS) test p-value for differences in the two distributions are indicated at the bottom of the class descriptions. E. CDF plot as in S3D for genes upregulated in *rbm8a* mutant embryos at 21 hpf (red) compared to the unchanged genes (black). F. CDF plot as in S3D for genes upregulated in *rbm8a* mutant embryos at 27 hpf (red) compared to the unchanged genes (black). G. Cumulative distribution frequency (CDF) plot showing the fold changes in *upf1* morphants (12 hpf) of genes upregulated in *rbm8a* mutant embryos at 21 hpf (red) compared to unchanged genes (black). Kolmogorov-Smirnov (KS) test p-value for differences in fold changes between the two groups is indicated on the bottom right.

**Figure S4. (related to Figure 6).**
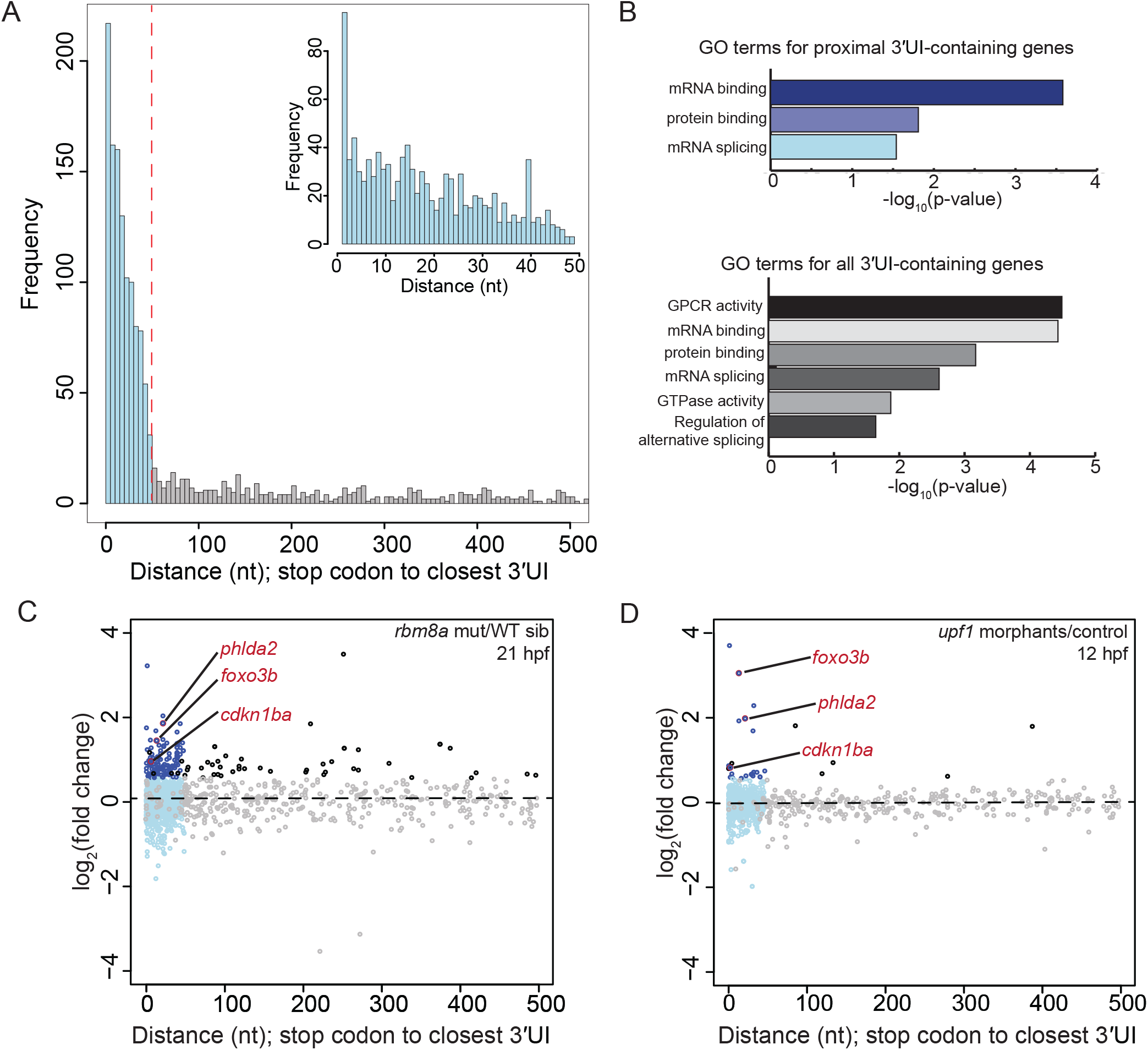
A. Histogram depicting the frequency of all zebrafish 3′UI transcripts in Ensembl GRCz10 as a measure of the distance of the 3′UI from the stop codon. Data are shown in 5 nts bins and bins beyond 500 nts are not shown. Bins of proximal 3′UI genes are in blue and distal 3′UI bins are in gray. Inset: Histogram of all zebrafish proximal 3′UI transcripts binned by 1 nt. B. PANTHER14.0 (70) gene ontology (GO) term enrichment analysis of proximal 3′UI-containing genes (top, shades of blue) and all 3′UI-containing genes (bottom, shades of gray). All significant terms (Benjamini-Hochberg corrected p-value < 0.05) are shown for each set. C. A scatter plot showing fold change (FC) for genes with proximal 3′UI (dark blue: FC > 1.5 and light blue: FC < 1.5) and distal 3′UI (black: FC > 1.5 and gray: FC < 1.5) in *rbm8a* mutant embryos at 21 hpf compared to wild-type siblings. Genes encircled in red also contain a proximal 3′UI in mouse and human, and were independently validated in (D). Gray and black dots in the < 50 nts region represent genes with one proximal and one or more distal intron. D. A scatter plot as in C showing fold changes of 3′UI-containing genes for 12 hpf *upf1* morphants compared to wild-type control embryos.

**Figure S5. (related to Figure 7).**
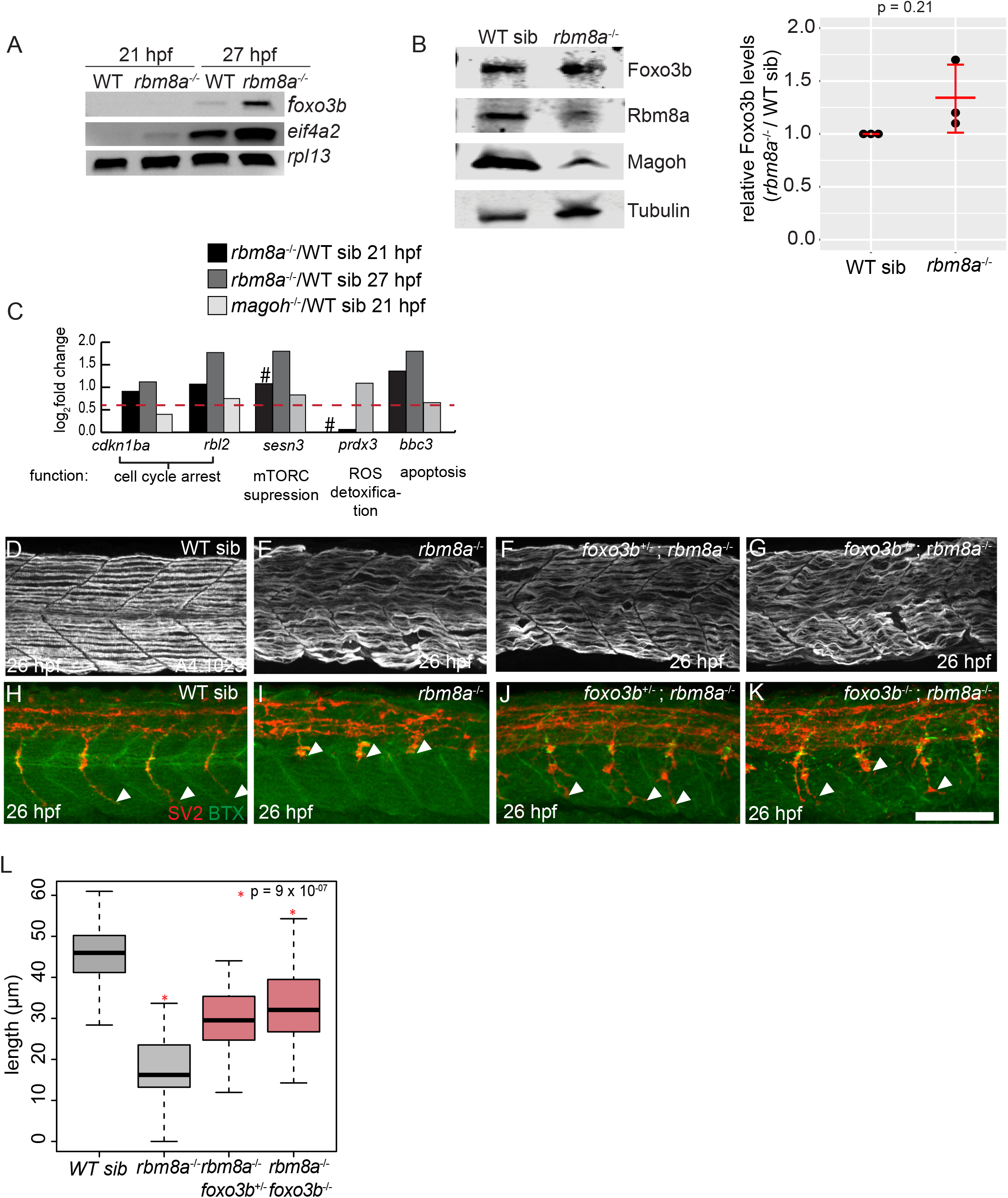
A. Semi-quantitative RT-PCR shows transcript levels of *foxo3b*, *eif4a2* and *rpl13* (loading control) in *rbm8a* mutant and wild-type sibling embryos at 21 and 27 hpf. B. Western blots (on the left) show levels of Foxo3b, Rbm8a, Magoh, and Tubulin in *rbm8a* mutant embryos compared to WT siblings at 27 hpf (N = 20 embryos per genotype). Right: dot plot showing Foxo3b levels normalized to tubulin levels in *rbm8a* mutant embryos and WT siblings at 21 hpf in three biological replicates. C. Bar graph showing log2 fold changes of known Foxo3b transcriptional targets that show a significant upregulation (FDR < 0.05) in EJC mutant RNA-Seq datasets. Foxo3b targets are from Morris et al. 2015 (46). A pound symbol indicates log2 fold change with FDR > 0.05. Horizontal dotted red line indicates fold change of 1.5. D-G. Confocal images showing Myh1 immunofluorescence using anti-A4.1025 in somites 12-16 of WT sibling (D), *rbm8a*^-/-^ mutant (E), *rbm8a*^-/-^; *foxo3b*^+/-^ mutant (F), and *rbm8a*^-/-^; *foxo3b*^-/-^ mutant (G) embryos. (N = 5 embryos/genotype). Scalebar in K (for panels D-K) is 100 nm. H-K. Merged confocal images showing motor neurons (red; detected by anti-SV2 staining) and acetylcholine receptors (green; detected by alpha-bungarotoxin staining) in somites 12-16 of WT sibling (H), *rbm8a*^-/-^ mutant (I), *rbm8a*^-/-^; *foxo3b*^+/-^ mutant (J), and *rbm8a*^-/-^; *foxo3b*^-/-^ mutant (K) embryos. Neuromuscular junctions in the merged image are yellow. White arrowheads point to the distal end of the motor neuron. (N = 5 embryos/genotype). Scalebar in K (for panels D-K) is 100 nm. L. Boxplots showing quantification of motor neuron length in embryos of genotypes indicated along the x-axis) (4 motor neurons/embryo and 5 embryos/genotype).

**Figure S6. (related to Figure 8).**
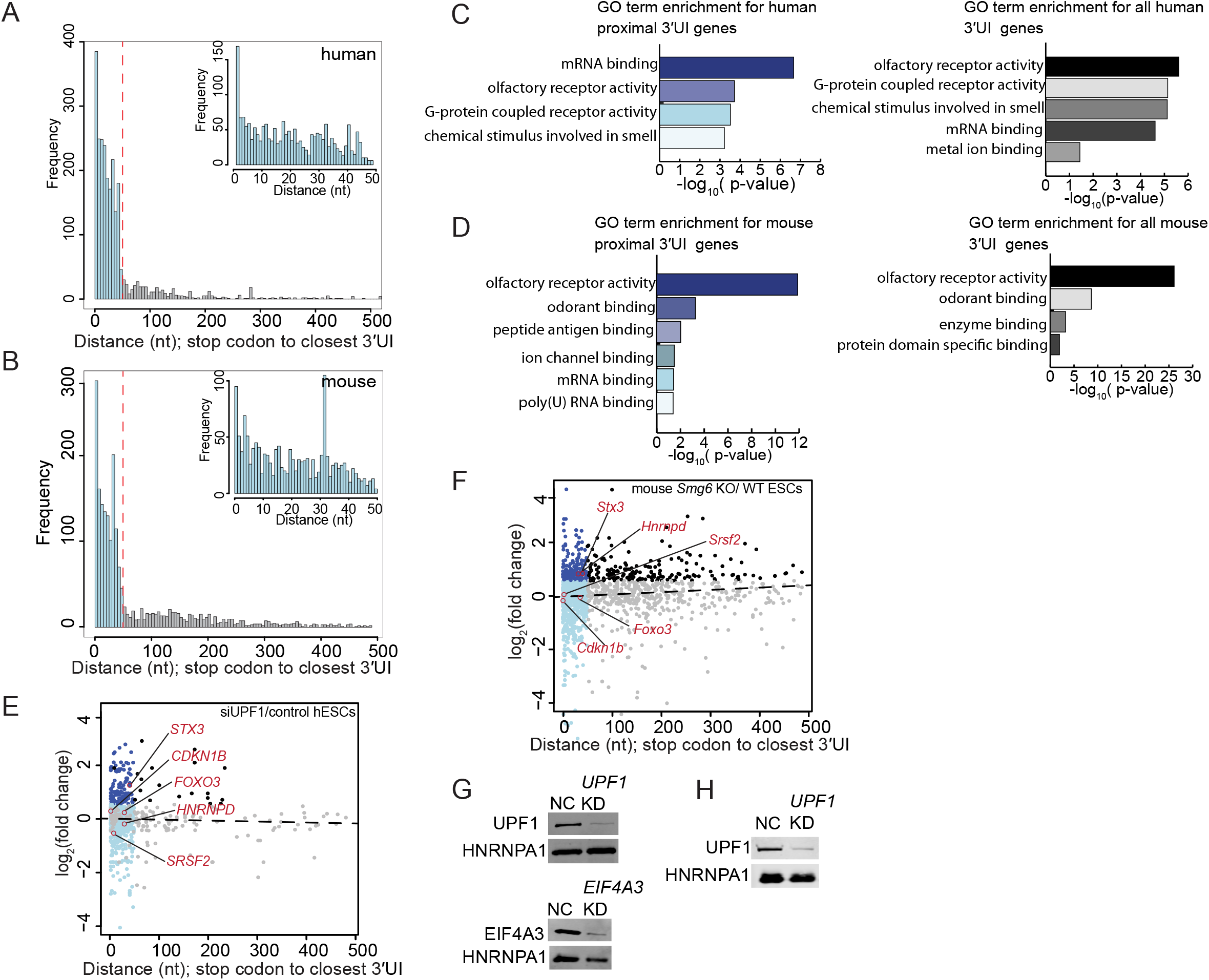
A. Histogram showing the frequency of all 3′UI-containing transcripts in human GRCh38 as a measure of the distance of the 3′UI from the stop codon. Data are grouped in 5 nt bins from 1-500 nts. Proximal 3′UI-containing gene bins are indicated in blue; distal 3′UI-containing gene bins are indicated in gray. Red dotted line indicates distance from stop codon to closest 3’UI = 50 nts. B. Histogram as in A of mouse proximal and distal 3′UI-containing genes in mouse GRCm38. C. PANTHER14.0 (70) gene ontology (GO) term enrichment analysis of proximal 3′UI-containing genes (shades of blue) and all 3′UI-containing genes (shades of gray). All significant terms (Benjamini-Hochberg corrected p-value < 0.05) are shown for each set. D. GO term enrichment analysis as in C of mouse 3′UI-containing genes. E. A scatter plot showing fold changes for all proximal 3′UI genes (dark blue: FC > 1.5 and light blue: FC < 1.5) and all distal 3′UI genes (black: FC > 1.5 and gray: FC < 1.5) in *UPF1* knockdown human embryonic stem cells (hESCs) compared to control cells using previously published data (data from 47). Gray and black dots in the < 50nts region represent genes with one proximal and one or more distal intron. F. Scatter plot as in E showing fold changes of mouse 3′UI-containing genes in *Smg6*^-/-^ knockout mouse embryonic stem cells (mESCs) compared to wild-type cells (51). G-H. Western blots showing representative protein knockdown in one replicate for Figures 8E (G) and 8F (H).

**Table S1**: RIP-Seq and RNA-Seq data summary

**Table S2**: Genes upregulated in RNA-Seq datasets represented in Fig 5A

**Table S3**: Master list of zebrafish, mouse and human 3′UI-containing genes

**Table S4**: List of oligonucleotides used in this study

## Materials and methods

### Animal stocks, lines, and husbandry

Adult zebrafish (*Danio rerio*) were housed at 28.5°C on a 14 hour light/10 hour dark cycle and embryos were obtained by natural spawning or *in vitro* fertilization. Embryos were raised at both 25°C and 28.5°C and were staged according to Kimmel et al. 1995 (71). *rbm8a*^oz35^ and *magoh*^oz36^ lines generated using CRISPR/Cas9 mutagenesis (described below) in the AB strain. The *foxo3b*^ihb404^ line (47) was obtained from the Xiao lab at the Chinese Academy of Sciences, Wuhan, China. Animal experiments were performed in accordance with institutional and national guidelines and regulations and were approved by the Ohio State University Animal Care and Use Committees.

### CRISPR/Cas9 mutagenesis

An optimal CRISPR target site in the coding sequence of *rbm8a* and *magoh* was identified using the ZiFit Targeter software package (72, 73). gRNAs were designed and synthesized as described (34). *rbm8a-* or *magoh*-targeting gRNA was co-injected with *Cas9* mRNA (74) into 1-cell stage embryos (60 pg gRNA and 160 pg *Cas9* mRNA). *rbm8a* gRNA target site: (5’-GGGAGGCGAAGACTTTCCTA-3’) *magoh* gRNA target site: (5′-GGTACTATGTGGGGCATAA-3′) Injected embryos were raised to 24 hpf at which time embryos were individually screened by high-resolution melting analysis (HRMA) to assess target site mutation efficiency in somatic cells. Remaining embryos were raised and crossed to AB wild-type adults; F1 adults were screened for germline transmission of CRISPR-induced mutations using HRMA. HRMA revealed unique *rbm8a* and *magoh* mutant alleles transmitted by multiple F0 founders. We recovered the *rbm8a^oz35^ and magoh^oz36^* alleles and outcrossed the heterozygotes to the AB wild-type strain for two generations before intercrossing for phenotypic analyses. Sequences of primers used for genotyping are listed in Supplemental Table 4.

### EJC mutant and *foxo3b^ihb404^* mutant embryo and adult genotyping strategy

Individual embryos and adult fin tissue were lysed in 50 µl 1M NaOH for 15 mins at 95°C followed by incubation on ice for 5 minutes at 4°C, and then neutralized with 5 µl of 1M Tris-HCl pH 8. For genotyping fixed embryos, heads were removed into ThermoPol buffer (20 µl) and treated with 2 mg/ml ProK at 55°C for 3 hours to extract DNA. 1 µl of DNA extract was used as a template in a 20 µl PCR with Taq polymerase according to the manufacturer’s protocol (NEB). For genotyping *rbm8a^oz35^ and foxo3b^ihb404^* mutant embryos, PCR products were digested with 20 units of XmnI and XcmI respectively (NEB) to distinguish cleavable mutant from un-cleavable wild-type amplicons. Digested products were analyzed on a 1% agarose gel stained with Gel Red (Biotium). For genotyping *magoh^oz36^* mutant embryos, PCR products were analyzed by separation of mutant and wild-type alleles on a 2% agarose gel stained with Gel Red (Biotium). Primer sequences are listed in Table S4.

### Acridine orange staining and immunohistochemistry

Embryos were incubated in 1:5000 acridine orange solution for 1 hr at 28.5°C (stock: 6 mg/ml, Sigma-Aldrich) followed by 2X washes in fish system water. For immunohistochemistry, embryos were processed following standard protocols using 4% PFA fixation, permeabilization using acetone, and incubation in blocking solution for 1 hour. EJC mutant embryos and wild-type siblings at 24 hpf and 26 hpf were incubated in 2% BSA/2% goat serum/1% DMSO/0.1% Tween-20/PBS blocking solution with 1:100 dilution anti-SV2 (DSHB) and 1:1000 anti-A4.1025 (DSHB) primary antibodies and AlexaFlour (Molecular Probes) secondary antibodies. Embryos were stained with Alexa Fluor 488-conjugated α-Bungarotoxin (Thermo Fisher) incubation in a 1:200 blocking solution post primary and secondary antibody staining. All images were centered on the region above the end of the yolk tube which included somites 12-16 at 24 hpf and somites 16-20 at 26 hpf.

### Microscopy and Imaging

Immuno-stained embryos were dissected and mounted in Fluoromount-G (SouthernBiotech) and imaged at 40x magnification using MetaMorph software (Molecular Devices) on an Andor™ SpinningDisc Confocal Microscope (Oxford Instruments) with Nikon Neo camera. Live images of EJC mutant and wild-type sibling embryos were taken by mounting embryos in 3% methylcellulose and imaging with a AxioCam camera on a Zeiss upright AxioPlan2 microscope.

### Zebrafish NMDI14 inhibitor treatment

NMDI14 (Sigma) stock solution was made in DMSO as per manufacturer’s instructions. AB wild-type zebrafish embryos were dechorionated on agarose-coated 10 cm plates. At 3 hpf, NMDI14 was added to a final concentration of 4.8 μM. At 24 hpf, embryos (20/treatment) were rinsed with fresh fish water and added to 500 μl of Trizol (Thermo Fisher Scientific) for RNA preparation.

### Immunoblot analysis

SDS-PAGE gels and western blots were performed using the standard mini-PROTEAN tetra system (Bio-Rad). All western blots were stained using infrared fluorophore-conjugated secondary antibodies and were scanned on a LI-COR Odyssey CLx imager. Protein quantification was performed using Image Studio software (v5.2.5).

### Quantification of paralysis and motor neuron length

At 24 hpf, EJC mutant and wild-type sibling embryo movements were scored under the dissecting microscope by counting the number of tail contractions per minute. For motor neuron length quantification, immunofluorescence images of 26 hpf EJC mutant and wild-type sibling embryos were stained with anti-SV2 as described above. Images were imported into Fiji (ImageJ v2) and motor neuron length was quantified using the Simple Neurite Tracer plugin (75).

### RNA-Immunoprecipitation-seq and RNA-Seq sample collection

At 24 hpf, zebrafish embryos (n= 800 embryos/IP) were triturated using a 200 µl pipette and washed to remove yolks as previously described (76), followed by flash freezing the tissue in liquid nitrogen. Whole embryo tissue was lysed and sonicated in 800 μl of hypotonic lysis buffer (HLB) [20 mM Tris-HCl pH 7.5, 15 mM NaCl, 10 mM EDTA, 0.5 % NP-40, 0.1 % Triton X-100, 1 mM Aprotinin, 1 mM Leupeptin, 1 mM Pepstatin, 1 mM PMSF]. Lysates were sonicated using a microtip for 7 seconds, NaCl was increased to 150 mM, and RNase I was added to 100 μg/ml. Following a 5-minute incubation on ice, cell lysates were cleared by centrifugation at 15,000 × g. The sample was split into 2 tubes (400 ul each) and the volume was increased to 2 mL by addition of isotonic lysis buffer. Complexes were captured on Protein G Dynabeads (Thermo Fisher) conjugated to IgG or α-Rbm8a for 2 hours at 4 °C. Complexes were washed in isotonic wash buffer (IsoWB) [20 mM Tris-HCl pH 7.5, 150 mM NaCl, 0.1 % NP-40] and eluted in clear sample buffer [100 mM Tris-HCl pH 6.8, 4 % SDS, 10 mM EDTA, 100 mM DTT]. The proteins were eluted in 20 μl of clear sample buffer [100 mM Tris-Hcl pH 6.8, 4% SDS, 10mM EDTA, 100 mM DTT] and 10 μl of the sample was used to separate the proteins via SDS-PAGE and analyze by western blotting. The remaining 10 μl of the sample was used for RNA extraction using Phenol-Chloroform-Isoamyl alcohol precipitation. RNA was resuspended in 10 μl of Rnase-free water. 1 μl of the RNA was end-labeled with γ- 32 ATP and then run on a denaturing 4% urea PAGE gel to assess quality while the remainder was used for RNA-seq library preparation.

For RNA-Seq sample collection, EJC mutant and wild-type sibling embryos (N = 25) were harvested at 21 and/or 27 hpf and lysed in 500 µl Trizol (Thermo Fisher Scientific). RNA was extracted following manufacturer standard procedures.

### RIP-seq and RNA-seq library preparation

For RIP-Seq, RNA extracted from ∼90 % of RIP eluate was used to generate strand-specific libraries. For RNA-Seq libraries, 5 µg of total cellular RNA was depleted of ribosomal RNA (RiboZero kit, Illumina), and subjected to base hydrolysis. RNA fragments were then used to generate strand-specific libraries using a custom library preparation method (77). Briefly, a pre-adenylated miR-Cat33 DNA adapter was ligated to RNA 3’-ends and used as a primer binding site for reverse-transcription (RT) using a special RT primer. This RT primer contains two sequences linked via a flexible PEG spacer. The DNA with a free 3’-end contains sequence complementary to a DNA adapter as well as Illumina PE 2.0 primers. The DNA with a free 5’-end contains Illumina PE 1.0 primer sequences followed by a random pentamer, a 5 nt barcode sequence, and ends in GG at the 5’-end. Following RT, the extended RT primer was gel purified, circularized using CircLigase (Illumina), and used for PCR amplification using Illumina PE 1.0 and PE 2.0 primers. All DNA libraries were quantified using an Agilent Bioanalyzer instrument to determine DNA length and a Qubit Fluorometer to quantify DNA amount. Libraries were sequenced on an Illumina HiSeq 2500 platform in the single-end format (50 nt read lengths).

### Zebrafish EJC mutant embryo RIP-seq and RNA-Seq data analysis

#### Adapter trimming and PCR duplicate removal

After demultiplexing, fastq files containing unmapped reads were first trimmed using Cutadapt (v2.3). A 12 nt sequence on read 5’-end consisting of a 5 nt random barcode sequence, 5 nt identifying barcode, and a CC was removed. The random barcode sequence associated with each read was saved for identifying PCR duplicates down the line. Next, as much of the 3’-adapter (miR-Cat22) sequence TGGAATTCTCGGGTGCCAAGG was removed from the 3’-end as possible. Any reads less than 20 nts in length after trimming were discarded.

#### Alignment and removal of multi-mapping reads

##### RIP-Seq

Following trimming, reads were aligned with HISAT2 v2.1.0 (Kim et al., 2015) using 24 threads to zebrafish GRCz10. After alignment, reads with a HISAT2 mapping score less than 60 were removed, i.e. all multi-mapped reads were discarded. Finally, all reads mapping to identical regions were compared for their random barcode sequence; if the random sequences matched, such reads were inferred as PCR duplicates and only one such read was kept.

##### RNA-Seq

Trimmed libraries were aligned to the zebrafish genome using TopHat2 (78) (v2.0.14 and default options: --read-mismatches 2, --red-gap-length 2, --read-edit-dist 2, --min-anchor-length 8, --splice-mismatches 0, --num-threads 2 (not default), --max- multihits 20) and the GRCz10 genome assembly. Read count followed by differential expression analysis was conducted as stated in the Love et al. 2018 RNA-Seq workflow.

#### RIP-Seq data downstream analyses

First, by comparing aligned reads to a GRCz10 exon annotation obtained through Ensembl BioMart we determined the 5’ and 3’ end distribution of RIP-Seq reads and the meta-exon distributions of RIP-Seq reads. The primary reference transcriptome was obtained from Ensembl BioMart. RIP-Seq transcripts with an APPRIS P1 annotation were filtered out and only one major transcript per gene was used for all analyses concerning the specificity of the RIP-Seq replicates. These analyses include calculation of RPKMs for the major APPRIS P1 isoform and compare intronic RPKMs to exonic RPKMs as well as intron-less transcript RPKMs to multi-exon transcript RPKMs.

#### RNA-Seq differential expression analysis

Differential expression analysis using EJC mutant embryos and wild-type RNA-Seq data was conducted based on the RNA-Seq workflow published by Love et al. 2018 (79). For each RNA-Seq experiment consisting of an EJC mutant and its WT sibling at a given time-point, three biological replicates were sequenced per genotype.

First, to create count-matrices for each RNA-Seq experiment, the GenomicAlignments and SummarizedExperiment software (80, 81) were used to count reads mapping per gene for each RNA-Seq bio-replicate. The count matrix was filtered to remove all genes with zero counts in all samples before differential expression (DE) analysis using DESeq2 (82). At least one, if not all of the biological replicates for each RNA-Seq experiment were sequenced during a separate deep-sequencing run. These differences in sequencing runs introduced some variability among the replicates. To account for the variability among our RNA-Seq bio-replicates during differential expression analysis we used the RUV-seq R package (83). Usual methods of normalization only account for sequencing depth but RUV-seq methods can be used to normalize libraries for library preparation and other technical effects. We used the RUVs method of the RUV-seq package which utilizes the centered counts (the counts of genes unaffected by our covariates of interest such as the sample genotype) to determine a normalization factor for each library. The count matrix was imported into DESeq2, and the RUVs normalization factors and genotype were used in the design formula to construct the DESeq dataset for gene-level differential expression analysis. We used the LRT test with all default DESeq2 settings to identify genes differentially expressed between mutant and wild-type samples. To correct for multiple testing in the DE analysis we used Benjamini-Hochberg (BH) adjustment with independentFiltering set to false. We decided to set independentFiltering to false because we are interested in studying NMD targets which are most likely to have low read counts in wild-type embryos. In the case of the *rbm8a*^-/-^ 21 hpf and 27 hpf datasets the histogram of all p-values showed a hill-shaped distribution. In order to account for this distribution, as per the suggestion made in RNA-Seq workflow (79), we used fdrtool (84) for multiple testing and determined adjusted p-values using default fdrtool settings.

### Gene Ontology enrichment analysis

The PANTHER14.0 (70) tool was used to identify significantly enriched biological process GO terms in genes that are found to be significantly differentially expressed in EJC mutant embryos by DESeq2. The PANTHER tool was also used to identify significantly enriched biological process GO terms in proximal 3′UI genes in zebrafish and humans. For all analyses the PANTHER’s Benjamini-Hochberg correction was used to calculate adjusted p-values.

### Overlap analysis

The universal and test sets were chosen to be the set of Ensembl gene IDs which did not correspond to NA in the adjusted p-value and gene symbol columns post-DESeq2 analysis. After determining a universal dataset for each RNA-seq dataset, the smallest universal set for the comparison in question was chosen. The significance of overlap was calculated using a hypergeometric test on the R statistical software.

### STRING network analysis

The STRING database (85) was used to identify connections between proteins encoded by proximal 3′UI genes with default high confidence settings (minimum required interaction score = 0.7). The clusters shown were created after the Markov Cluster Algorithm (MCL) inflation parameter was set to 3 clusters.

### Human and mouse RNA-Seq data analysis

SRA files were downloaded from sources specified in (50, 51). Fastq files generated from the SRA files were mapped using TopHat2 (version 2.1.1) using the same settings described above for zebrafish alignment. Count matrices were generated using the GenomicAlignments and SummarizedExperiment packages. The count matrices were imported into DESeq2 for differential expression analysis using the LRT test, BH adjustment and with independentFiltering set to false.

### Identification of uORF genes in zebrafish

We selected for uORFs which were categorized as “functional uORFs” in Johnstone et al. 2016 (41) based on RNA-Seq and ribosome profiling. The following filters were applied to select for uORFs that show evidence of translation at 24 hpf: 5’UTR RPF RPKM ≥ 5, RNA-Seq RPKM≥ 5, RPF RPKM ≥ 5, ORF translation efficiency at 24 hpf >1.

### Identification of 3′UTR intron containing genes in zebrafish, mouse, and human

A table describing exon starts, exon ends, CDS start, CDS end, strand and APPRIS annotation was downloaded from the Ensembl database for all transcripts in zebrafish (GRCz10), human (GRCh38) and mouse (GRCm38). All transcripts with any level of APPRIS annotation (39) were included. We then identified transcripts that contain introns in the 3′UTRs by subtracting exon start coordinates from the CDS end coordinates in a strand specific manner. We then determined the distance of the nearest 3′UTR intron to the stop codon; based on the distance (< or ≥ 50 nts) as well as the number of 3′UTR introns the transcripts were classified into proximal and distal categories. Proximal transcripts were defined by the presence of only one 3′UTR intron which is within 50 nts of the normal stop codon. Distal transcripts were defined by the presence of one 3′UTR intron which is more than 50 nts away from the stop codon or by the presence of more than one 3′UTR intron irrespective of the distance of the nearest 3′UTR intron to the stop. For all fold-change analyses, we defined distal/proximal 3′UI-containing genes as those that encode one or more distal/proximal 3UI+ transcripts. The distal 3UI-containing genes that did not have an APPRIS annotation but were annotated with an ‘NMD biotype’ in the Ensembl database were included as a separate group in the analyses included in Fig. 5 and the group was named as NMD biotype.

### Mammalian cell culture knockdown experiments

HCT116 cells were seeded into 12 well plates (10^5^ cells/ well) in McCoy’s 5A media and 15 pmol siRNA was reverse transfected using 1.6 µl lipofectamine RNAiMAX reagent (Thermo Fisher Scientific) per well. Knockdown was carried out for 48 hours with a media change after 24 hours. Cells were harvested in hypotonic lysis buffer (described above), 30% of the cell lysate was saved to check the efficiency of knockdown while the rest was added to TRI Reagent (Sigma) for RNA extraction. The siRNAs used in this study are listed below:

Hs_*RBM8A*_5 FlexiTube siRNA, no modification, 20 nmole (SI03046533, Qiagen) All Stars Negative Control siRNA, no modification, 20 nmole (SI03650318, Qiagen)

*UPF1*_1879: AAG AUG CAG UUC CGC UCC AUU

*EIF4A3*_187: CGA GCA AUC AAG CAG AUC AUU

### Zebrafish *upf1* knockdown experiment and RNA-Seq

For conducting *upf1* knockdown in zebrafish, 2 ng of a splice blocking morpholino (MO) diluted in 0.2 M KCl with 0.1% phenol red was injected into 1-cell stage embryos. The *upf1* MO used was previously published and named *upf1* MO2 (37) with a sequence of 5’-TTTTGGGAGTTTATACCTGGTTGTC-3’. Morpholino was synthesized by Gene Tools, LLC. Uninjected wild-type control embryos (n=30) and injected morphants (n=30) were raised for 12 hours at 28.5°C and lysed in 500 µl Trizol following manufacturer’s procedures (Thermo Fisher Scientific). After RNA-extraction, double-stranded cDNA was synthesized following Illumina’s TruSeq protocol per manufacturer’s instructions. Briefly, mRNA was purified from 1 µg total RNA using Dynabeads oligo(dT)25 magnetic beads (Thermo Fisher Scientific) followed by clean-up with AMPure XP SPRI beads (Beckman Coulter). mRNA was then fragmented for 5 minutes at 70°C using Ambion’s RNA Fragmentation Reagent (AM8740) followed by an additional clean-up step using AMPure XP SPRI beads. Fragmented RNA was reverse transcribed using random primers and SuperScript III (Thermo Fisher Scientific). After second strand synthesis, end repair, and 3’ end adenylation per the TruSeq protocol (Illumina, Inc.), libraries were constructed using an Apollo 324 automated library system. After Illumina adapter ligation and amplification, all DNA libraries were quantified using an Agilent Bioanalyzer instrument to determine DNA length and a Qubit Fluorometer to quantify DNA amount. 10 cycles of amplification was performed prior to sequencing on an Illumina HiSeq 2000 system in the paired-end format (100 nt read lengths). All experiments were performed in biological duplicate.

#### RNA-seq data analysis

Trimmed reads were obtained from the sequencing core and then mapped to the GRCz10 genome assembly using TopHat2 as described above. Differential gene expression analysis was also performed as described above using DESeq2. RUV-Seq and fdrtool corrections were not required.

### Quantitative RT-PCR

Zebrafish embryos and mammalian cells were harvested in Trizol. RNA was isolated using standard Trizol procedures, followed by DNase treatment, purification with Phenol:Chloroform: Isoamyl alcohol (25:24:1, pH 4.5) and resuspension in RNase-free water. 1.5 μg of RNA was reverse transcribed using oligo-dT and Superscript III (Invitrogen). After reverse transcription of RNA, the samples were treated with RNase H (Promega) for 30 min at 37°C. For each qPCR 30 ng of cDNA was mixed with 5 μl of 2X SYBR Green Master Mix (ABS), 0.2 μl of a 10 mM forward and reverse primer each (defrosted once) in a 10 μl reaction. The qPCRs were performed in triplicate (technical) using primers described in Table 4. Reference genes for relative quantification were *mob4* in zebrafish (86) and TATA-binding protein (*TBP*) in human cells. Fold-change calculations were performed by the ΔΔCt method. Fold-changes from three biological replicates were used to determine the standard error of means. The p-values were calculated using Welch t-test in the R statistical computing software.

### NMD reporter constructs and Northern blotting

3′UTR sequences for *ARC, FOXO3* and *HNRNPD* were amplified from HeLa cell genomic DNA (for 3′UI+ constructs) or cDNA libraries (for 3′UI-constructs) and inserted into NotI and XbaI sites of pcTET2-βwtβ (gift from Jens Lykke-Andersen). For the *FOXO3* 3′UI+ construct, only ∼250 nts at the start and end of the 3′UI was included to create a “mini” intron. For analysis of mRNA steady state levels, the 3′UI+ or 3′UI-reporters were co-transfected along with a control *β-globin* expressing plasmid (pcβwtβ or pcβwtGAP3UAC, both gifts from JLA). To knockdown UPF1, 15 pmol of siRNA is reverse transfected to 7×10^4^ HeLa-Tetoff cells (Clontech) in a 12 well plate using 1.6 µl RNAiMAX (Thermo Fisher Scientific) and 200 µl Opti-MEM (Gibco). After 18 hours, plasmids (150-250 ng 3’UI constructs, 20-30 ng pcβwtGAP3UAC or pcβwtβ and 20-30 ng pezYFP) were co-transfected with the second dose of 15 pmol siRNA with 0.9 µl JetPrime transfection reagent (Polyplus). Tetracycline was added to 100 ng/ml to inhibit the expression of the 3’UI reporter. 40 hours after the reverse transfection, expression of the 3’UI reporter was induced by removing tetracycline and incubation for an additional 8 hours, followed by cell harvesting and addition of either hypotonic lysis buffer (described above) for western blotting or TRI Reagent (Sigma) for RNA isolation and northern blotting. RNA was extracted and analyzed via northern blotting as described previously in Mabin et al. 2018 (33).

### Quantification and statistical analysis

All western blots were performed using infrared fluorophore conjugated secondary antibodies and were scanned on a LI-COR Odyssey CLx imager. Protein quantification was performed using Image Studio software (v5.2.5). Northern blot autoradiograms were scanned using Fuji FLA imager and quantified using ImageQuant TL software (v7.0). Average and standard error of means in the observed signal was determined for data from at least three biological replicates.

## Aknowledgements

We thank Pearlly Yan, the OSU Comprehensive Cancer Center genomics core and the Functional Genomics Laboratory at UC Berkeley for sequencing, Javier Cáceres and Dasa Longman for the zebrafish microarray data, Jens Lykke-Andersen for plasmids and antibodies, Xiao lab for zebrafish *foxo3b* CRISPR mutant line, Can Cenik, Christine Beattie, Robin Wharton and Anita Hopper for advice and critical comments. We also thank Caleb Embree, Anupama Rao, Derek Boehm and Jake Panten for their contributions to the project. This work was supported in part by an allocation from the Ohio Supercomputer Center. Funding for this work was provided by the Ohio State University, NIH/NINDS grant R01NS098780 (to S.L.A.), NIH/NIGMS grant R01GM117964 (to S.L.A.), NIH/NIGMS grant R01GM120209 (to G.S.) and the OSU Center for RNA Biology by the means of a seed grant to G.S. and S.L.A., graduate student fellowship to P.G.,Z.Y and summer fellowship to M.A.P. Instrumentation and core services were supported by the NIH grants S10-OD023582, P30-NS045758, P30-NS104177, and S10-OD010383.

## Author contributions

Conceptualization, P.G., G.S. and S.L.A; Investigation, P.G., T.L.G., R.D.P., Z.Y., K.T.T., M.A.P, N.C.D, S.L.A and G.S.; Writing – Original Draft, P.G. and G.S.; Writing – Review & Editing, P.G., T.L.G., R.D.P., Z.Y., K.T.T., M.A.P, N.C.D, R.B., S.L.A. and G.S.; Funding Acquisition, S.L.A. and G.S.; Resources, S.L.A. and G.S; Supervision, S.L.A. and G.S., and R.B.

## Data availability

All short read sequencing data will be submitted to and available at NCBI GEO database.

